# Visual cue-related activity of cells in the medial entorhinal cortex during navigation in virtual reality

**DOI:** 10.1101/453787

**Authors:** Amina A. Kinkhabwala, Yi Gu, Dmitriy Aronov, David W. Tank

## Abstract

During spatial navigation, animals use self-motion to estimate positions through path integration. However, estimation errors accumulate over time and it is unclear how they are corrected. Here we report a new cell class (“cue cell”) in mouse medial entorhinal cortex (MEC) that encoded visual cue information that could be used to correct errors in path integration. Cue cells accounted for a large fraction of unidentified MEC cells. They exhibited firing fields only near visual cues during virtual navigation and spatially stable activity during navigation in a real arena. Cue cells’ responses occurred in sequences repeated at each cue and were likely driven by visual inputs. In layers 2/3 of the MEC, cue cells formed clusters. Anatomically adjacent cue cells responded similarly to cues. These cue cell properties demonstrate that the MEC circuits contain a code representing spatial landmarks that could play a significant role in error correction during path integration.

## Introduction

Animals navigate using landmarks, objects or features that provide sensory cues, to estimate spatial location. When sensory cues defining position are either absent or unreliable during navigation, many animals can use self-motion to update internal representations of location through path integration (Mittelstaedt, 1982; Tsoar et al., 2011). A set of interacting brain regions, including the entorhinal cortex, parietal cortex, and the hippocampus (Brun et al., 2008; Bush et al., 2015; Calton et al., 2003; Calton et al., 2008; Clark et al., 2010; Clark et al., 2013; Clark et al., 2009; Clark and Taube, 2009; Frohardt et al., 2006; Geva-Sagiv et al., 2015; Golob and Taube, 1999; Golob et al., 1998; Hollup et al., 2001; Moser et al., 1993; Parron et al., 2004; Parron and Save, 2004; Taube et al., 1992; Whitlock et al., 2008) participate in this process.

The MEC is of particular interest in path integration. Grid cells in the MEC have multiple firing fields arrayed in a triangular lattice that tiles an environment (Hafting et al., 2005). This firing pattern is observed across different environments, and the grid cell population activity coherently shifts during locomotion (Fyhn et al., 2007). These observations have led to the hypothesis that grid cells form a spatial metric used by a path integrator. Given this, theoretical studies have demonstrated how velocity-encoding inputs to grid cell circuits could shift grid cell firing patterns, as expected of a path integrator (Barry and Burgess, 2014; Burak and Fiete, 2009; Fuhs and Touretzky, 2006; McNaughton et al., 2006). Cells encoding the speed of locomotion have been identified in this region (Kropff et al., 2015), providing evidence of velocity-encoding inputs and providing further support for the role of MEC in path integration.

A general problem with path integration is the accumulation of errors over time. A solution to this problem is to use reliable spatial cues to correct estimates of position (Evans et al., 2016; Hardcastle et al., 2015; Pollock, 2018). Many recent experimental studies showed profound impairment of grid cell activity by altering spatial cues, including landmarks and environmental boundaries. For example, the absence of visual landmarks significantly disrupted grid cell firing patterns (Chen et al., 2016; Perez-Escobar et al., 2016). Experiments that maintained the boundaries of a one-dimensional environment but manipulated nonmetric visual cues caused rate changes in grid cells (Perez-Escobar et al., 2016). The decoupling of an animal’s self-motion and visual scene altered grid cell firing patterns (Campbell et al., 2018). Also, many studies have also shown that grid cell firing patterns were also influenced by nearby boundaries (Carpenter et al., 2015; Derdikman et al., 2009; Giocomo, 2016; Hardcastle et al., 2015; Krupic et al., 2015; Krupic et al., 2018; Stensola et al., 2015; Yamahachi et al., 2013).

Border cells in the MEC, with firing fields extending across environmental boundaries (Solstad et al., 2008), are good candidates for supplying information for error correction near the perimeter of simple arenas (Pollock, 2018). This role of border cells is supported by the fact that an animal’s interactions with boundaries yielded direction-dependent error correction (Hardcastle et al., 2015). However, grid cell firing fields are maintained throughout open arenas in locations where border cells are not active and thus cannot participate in error correction. It is unclear how grid cells error correct to maintain stable firing fields within open areas of a bounded arena.

Also, natural navigation involves moving through landmark-rich environments with higher complexity than arenas with simple boundaries. How information from a landmark-rich environment is represented within the MEC is unknown. If there were cells in the MEC that encoded sensory information of landmarks, then more robust path integration and error correction of grid cells would be possible using circuitry self-contained within this brain area. In the MEC, while border cells have been shown to respond to landmarks in virtual reality (Campbell et al., 2018), increasing evidence suggests that unclassified cells also contained information about spatial environments (Diehl et al., 2017; Hardcastle et al., 2017; Hoydal, 2018; Kinkhabwala, 2015). Therefore, it would be useful to further determine whether these unclassified cells represent spatial cues (other than borders) that could be used in error correction.

Here we addressed this question by recording from populations of cells in the MEC during virtual navigation along landmark-rich linear tracks using electrophysiological and two-photon imaging approaches. Virtual reality (VR) allowed for complete control over the spatial information of the environment, including the presence of spatial cues along the track. The animal’s orientation within the environment was also controlled, simplifying analysis. We report that a significant fraction of the previously unclassified cells in MEC respond reliably to prominent spatial cues. As a population, the cells fire in a sequence as a spatial cue was passed. They were also anatomically organized by their spatial firing patterns within layers 2/3 of the MEC. These cells could provide the information necessary in local MEC circuits for error correction during path integration in sensory rich environments that are regularly found in nature.

## Results

### Cue-responsive cells in virtual reality

Mice were trained to unidirectionally navigate along linear tracks in virtual reality to receive water rewards. Virtual tracks were 8 meters long and had a similar general organization and appearance: the tracks began with a set of black walls, followed by a short segment with patterned walls, and then ended with a long corridor with a simple wall pattern (Figure 1A). Different environments were defined by different pairs of identical visual cues (tower-like structures) present along both sides of the corridor. These cues were non-uniformly spaced along the track. The last cue was always associated with a water reward.

**Figure 1.**
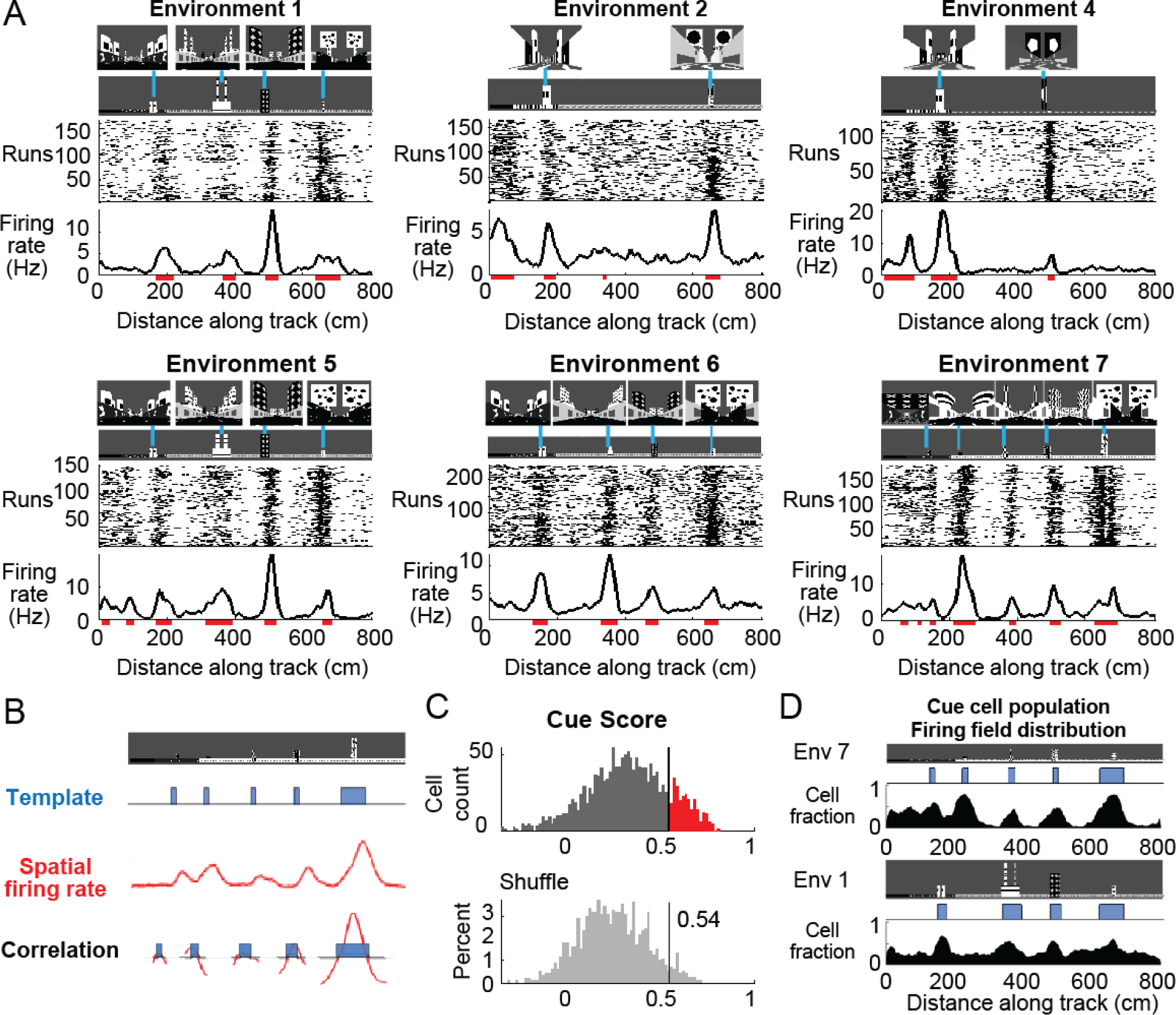
Cells respond to cues in the environment. A. Examples of cells with cue-related activity recorded during navigation along virtual tracks. At the top of each example are views of each cue from the animal’s perspective inside the track at that location. Side views of the track are shown below, with the start location to the left. The raster plot for a single cell’s spatial activity pattern across multiple traversals of the track is plotted with the average firing rate (Hz) as a function of track position (spatial firing rate) below. Spatial firing fields for the cell are indicated with horizontal red bars. B. Calculation of cue score. The Pearson correlation between the cue template and the cell’s spatial firing rate was calculated, then the cue template was shifted to where it was maximally correlated with the spatial firing rate and the correlations of cue template and spatial firing rate at each cue were individually calculated. The cue score was defined to be the mean of these correlations (Materials and Methods). C. The distribution of cue scores of recorded cells is shown at the top with the distribution of cue scores calculated on shuffled data shown below. The threshold was chosen as the value that 95% of the shuffled scores did not exceed (vertical black line). Cells exceeding this threshold were termed ‘cue cells’ and are shown in red in the top plot. D. Distribution of spatial firing fields of all cue cells in two environments.

We used tetrodes to record 1590 units in the MEC of four mice (Materials and Methods and Supplementary Figure 1). Activity of a subpopulation of these units exhibited a striking pattern, with spiking occurring only near cue locations along the virtual linear tracks (Figure 1A). On each run along the track, clusters of spikes were present at cue locations, forming a vertical band of spikes at each cue in the run-by-run raster plot. Spatial firing rates were calculated by averaging this spiking activity across all runs along the track. Clear peaks in the spatial firing rate were present at cue locations. We also defined spatial firing fields as the locations along the track where the spatial firing rate exceeded 70% of the shuffled data (Materials and Methods) and observed that the spatial firing fields were preferentially located near cue locations.

To quantify this feature of the spatial firing rate, we developed a “cue score” that measures the relationship between a cell’s spatial firing rate and the visual cues of the environment (Figure 1B and Materials and Methods). The cue score was based on the correlation of the cell’s spatial tuning with a spatial template that had value one at each cue, and zero elsewhere. Cells with cue scores above the threshold (95^th^ percentile of shuffled data, Materials and Methods) represented ~18% of all recorded cells (Figure 1C). In the remainder of the paper, we refer to these cells as “cue cells”.

We next quantified the distribution of spatial firing fields of all cue cells along the track by calculating, for each 5 cm bin, the fraction of cue cells with a spatial firing field (Materials and Methods). We defined the plot of this fraction versus location as the field distribution for all cue cells. This field distribution had peaks in locations where salient information about the environment was present (Figure 1D), and some fields were correlated with the beginning of the track where wall patterns changed. The mean firing field fraction for spatial bins in cue regions (0.4 ± 0.2) was higher than that for bins outside of cue regions (0.2 ± 0.1) (paired one-tailed t-test: cell fraction in cue regions > cell fraction in outside cue regions, N = 283, p < 0.001). Thus, the cue score identifies a subpopulation of MEC cells with spatial firing fields correlated with prominent spatial landmarks.

### Cue cell responses to environment perturbations

Is the activity of cue cells truly driven by the visual cues of the environment? To address this question, we designed related pairs of virtual tracks. One track had all cues present (*with-cues* track) and, in the second track, the last three cues were removed (*missing-cues* track). Tetrode recordings were performed as mice ran along both types of track in blocks of trials within the same session. Water rewards were delivered in the same location on each track regardless of the cue location differences.

At the beginning of both *with-cues* and *missing-cues* tracks where the tracks were identical, the spatial firing rates of cue cells were similar across tracks. Vertical bands of spikes were present in the run-by-run raster plots of both tracks and formed peaks in the spatial firing rate. The bands were also identified as spatial firing fields, which generally aligned to features of the environment (spatial cues/changes in wall patterns) present on both tracks (Figure 2A). However, the firing patterns changed dramatically from the point along the track where the environments began to differ. Spatial firing fields were prominent at cue locations along the entire remaining part of the *with-cues* track (Figure 2A, top) but were not present on the same part of the *missing-cues* track (Figure 2A, bottom). To quantify this difference, we compared changes in both cue cell spatial firing rates (Figure 2B) and in the spatial firing field distribution of the cue cell population (Figure 2C) across the two tracks. The data were split into two regions along the track (Figure 2B, left): the start region where cues were present for both tracks (gray bar marks this region, Region A - same) and the rest of the track where cues were either present or absent (blue bar, Region B - different). While cue cells showed similar mean and maximum firing rates in Region A across tracks, in Region B, mean firing rates remained the same but the maximum firing rates were lower in the *missing-cues* track compared to the *with-cues* track (maximum firing rate *with-cues*: 8.9 ± 5 Hz, *missing-cues:* 7.4 ± 4.9 Hz, mean difference (*with-cues* – *missing-cues*): 1.5 Hz; paired one-tailed t-test: maximum firing rate in Region B on *with-cues* track > maximum firing rate in Region B on *missing-cues* track, N = 161, p < 0.001). Moreover, cue cells had spatial firing fields clustered in each region where a cue was located on both tracks, but these fields were not present when cues were removed in Region B of the *missing-cues* track (Figure 2C). The mean fraction of cells with spatial firing fields was significantly higher when cues were present in comparison to when they were removed (Region B in cue bins: mean cell fraction on *with-cues* track: 0.41 ± 0.01, on *missing-cues* track: 0.33 ± 0.04; paired one-tailed t-test: *with-cues* cell fraction > *missing-cues* cell fraction, N = 161, p < 0.02). These results demonstrate that cue cells are more coherently active in regions along an environment where cues are located. When these cues are removed, these cells remain active but they do not coherently form spatial firing fields, indicating that the responses of these cells are correlated to the presence of the cues.

**Figure 2.**
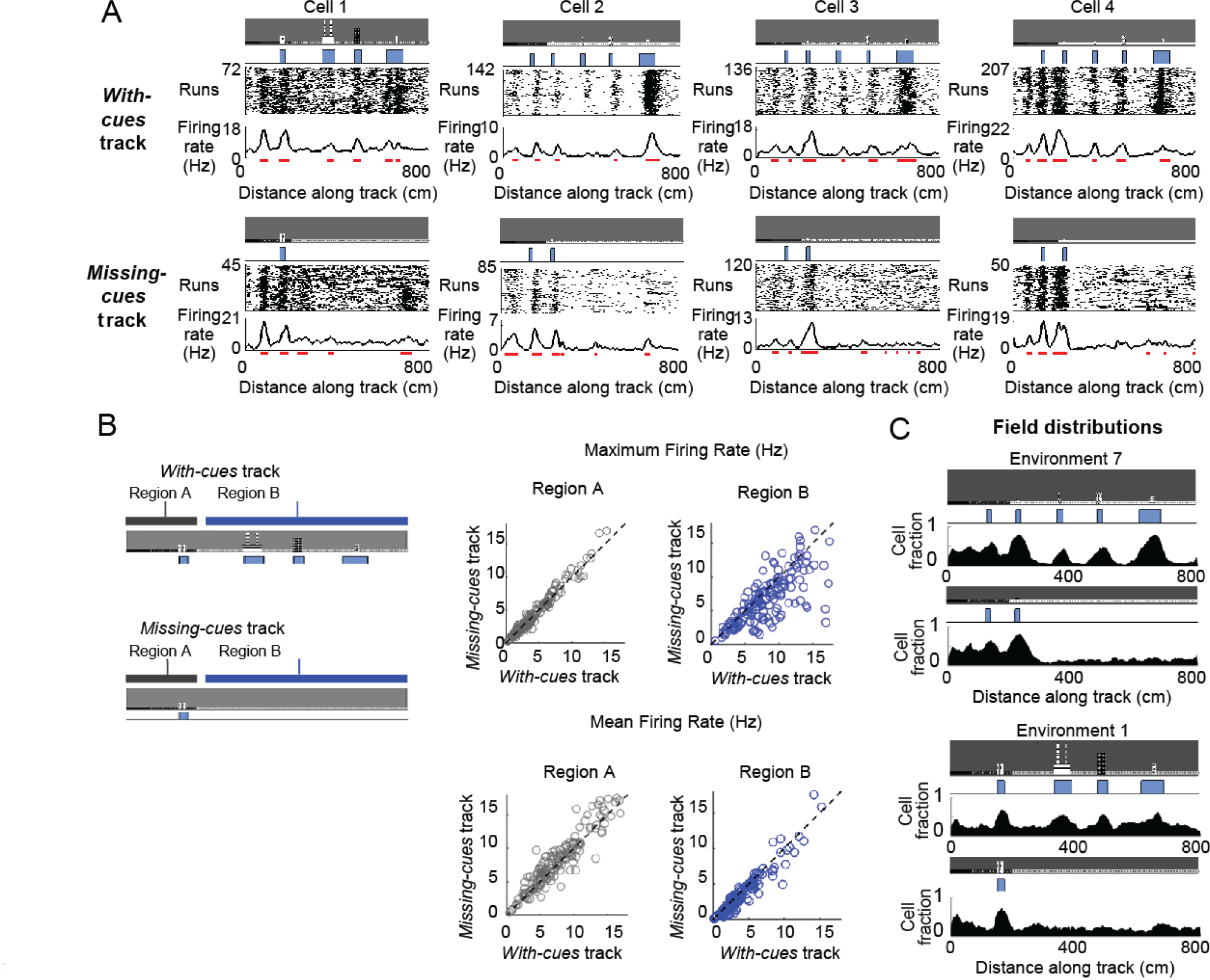
Cells respond to cue changes in environment. A. Examples of the spatial firing rates of cells during cue perturbation experiments. For each example, the top and bottom panels are from the same cell in blocks of trials in which the animal either ran down a virtual track with all cues present (*with-cues* track, top) or a track where some cues were missing in the later part of the track (*missing-cues* track, bottom). The environment and cue template for both environments are shown with the corresponding raster plots and spatial firing rates below. Cell 1: T5 20140211 t7 c2, Cell 2: T8 20140315 t8 c2, Cell 3: T10 20140326 t2wref c2, Cell 4: T10 20140313 t6wref c3. B. Comparison of firing rates of all cue cells between runs in the initial region that is the same for both tracks (Region A) and the later region (Region B) that either had cues (*with-cues* track) or missed cues (*missing-cues* track) (left). The maximum and mean firing rates in these regions of the two tracks are plotted to the top and bottom on the right. C. Population field distribution for the entire population of cue cells along *with-cues* and *missing-cues* tracks for two environments.

### Relationship to previously defined cell classes

To relate our population of cells recorded along linear tracks in virtual reality to previously characterized cell types in the MEC, the same cells were also recorded as the animal foraged for chocolate chunks in a real two-dimensional (2D) environment (0.5m × 0.5m). From the recordings in the real arena, we calculated grid, border, and head direction scores for all recorded cells (i.e. both cue and non-cue cells, Materials and Methods). We plotted the values of these spatial scores against the cue scores, which were calculated for the same cell during VR navigation, to determine the relationship of cue cells and the previously defined cell classes (Figure 3A). We found that a small percentage of cue cells were conjunctive with border (10%) or grid (18%) cell types, but the majority of cue cells had a significant head direction score (53%) (since the head direction score is based on orientation tuning, we do not consider the head direction cell type to be a spatial cell type). The total percentage of cue cells (18%) in the dataset was comparable to that of grid and border cells (Figure 3B).

**Figure 3.**
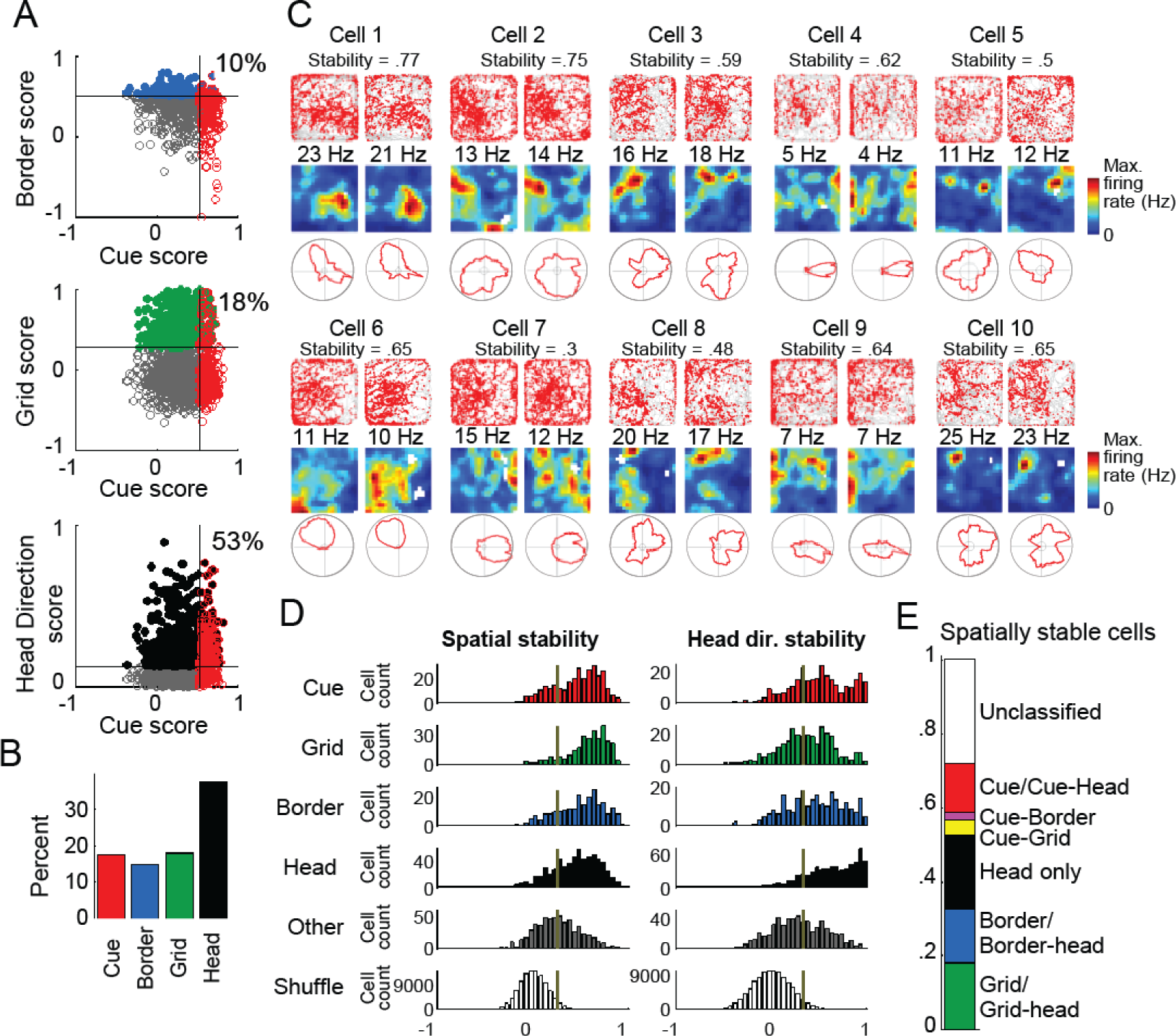
Cue cell activity during foraging in a real arena. A. Relative distributions of cue scores compared to border, grid, and head direction scores. Thresholds were calculated as the value that exceeds 95% of shuffled scores. The solid line indicates the threshold for each score that was used to determine the corresponding cell type (Materials and Methods). Cells are color-coded for whether they are cue (red), grid (green), border (blue), or head direction cells (black). The percentage of the cue cell population that was conjunctive for border, grid, and head direction is shown on each plot. B. Percentage of each cell type in the dataset. C. Examples of cue cells’ spatial firing rate in a real arena and their spatial stability. The recording of each cell was divided in half. The activity features of the first and second halves are shown for each cell in the left and right columns. For each column of each cell: top: plots of spike locations (red dots) and trajectory (gray lines); middle: the 2D spatial firing rate (represented in a heat map) with the maximum firing rate indicated above; bottom: head direction firing rate. The stability was calculated as the correlation of these two firing rates and shown under the cell number. D. Histograms of the spatial and head direction stability of the 2D real environment firing rates by cell type. E. Percentage of 2D real environment stable cells that are of a certain type. Cell types are color-coded: red = cue cell, green = grid cell, blue = border cell, black = head direction cell. The threshold for spatial stability was calculated as the value that exceeds 95% of score values for the shuffled distribution.

Since most cue cells (72%) were not conjunctive with a previously known spatial cell type, we next examined what their spatial activity patterns were in the real arena. As expected from their scores, most cells had irregular activity patterns in the arena and were not classified as any previously identified spatial cell type (Figure 3C and Supplementary Figure 2, the activity patterns of cue cells in the real environment are shown in the top and middle panels).

One striking feature of the spatial firing patterns of cue cells observed in real environments was the spatial stability of these complex and irregular patterns (Materials and Methods). The spatial firing rates in the real arena from the first and the second halves of the recording for 10 cue cells are shown in Figure 3C. We calculated the spatial stability as the correlation between these two halves and found that the spatial firing patterns were irregular but surprisingly stable. To further quantify this observation, we calculated the distributions of the stability of both the spatial and head direction firing rates for all cell types (Materials and Methods) (Boccara et al., 2010). We found that the distributions of stability for unclassified cells (not classified as cue, grid, border, or head direction cells, labeled as “Other”) were generally shifted towards lower values compared to all the currently classified cells, indicating that a large fraction of the remaining unclassified cells do not stably encode spatial and head direction information in the real arena (Figure 3D). Figure 3E shows the fractions of all cells with significant stability scores for their spatial firing rates in the real arena, classified by cell type. While some cue cells were conjunctive with border or grid cells, a large percentage (72%) of cue cells were previously unclassified as a particular spatial cell type. Cue cells accounted for 13% of the population of spatially stable cells, and for 22% of the previously unclassified spatially stable cells.

### Cue cells form a sequence at cue locations

For many cue cells, we observed that their firing fields had varying spatial shifts relative to the cue locations on virtual tracks (Figure 1A). We next investigated whether these shifts were consistent for a given cell across cues, suggesting that the spatial shift is a property for each cue cell similar to the spatial shift that defines the location of grid cell firing patterns. This would provide an overall patterning of the population of cue cell activity within an environment.

To test this, we took all cue cells identified for each virtual track and ordered their spatial firing rates and fields by the values of their spatial shift relative to the cue template, which was the smallest displacement of the cue template to best align with the firing rate (Figures 4A and 4B; Materials and Methods). We found a striking pattern where cue cells formed a sequence of spatial firing fields that was repeated at each cue. To examine if this pattern was produced by the concentration of neural firing around cues, rather than the alignment and ordering of the data alone, we compared this pattern to that of time-shuffled data, which were created by circularly permuting spike times of each cell by a random amount of time (Supplementary Figure 3, Materials and Methods). Time shuffled data did not exhibit an obvious sequence. This difference between the cue cells and shuffled data was further quantified by a ridge-to-background ratio (Materials and Methods), which was computed as the mean firing rate in a band centered on the sequential spatial firing rates of the cue cell population divided by the mean background rate outside of this band. We note that although ordering the spatial firing rates of the cells by their spatial shift was expected to create a ridge of firing rate along the diagonal, the mean ridge/background ratio for cue cells (2.58) was higher than that for time shuffled data (1.04 ± 0.003, p <0.001; N = 283 cue cells, 1,000 shuffles). Thus, the sequence represents sequential neural activity preferentially located near cue locations, rather than an artifact of ordering the data.

**Figure 4.**
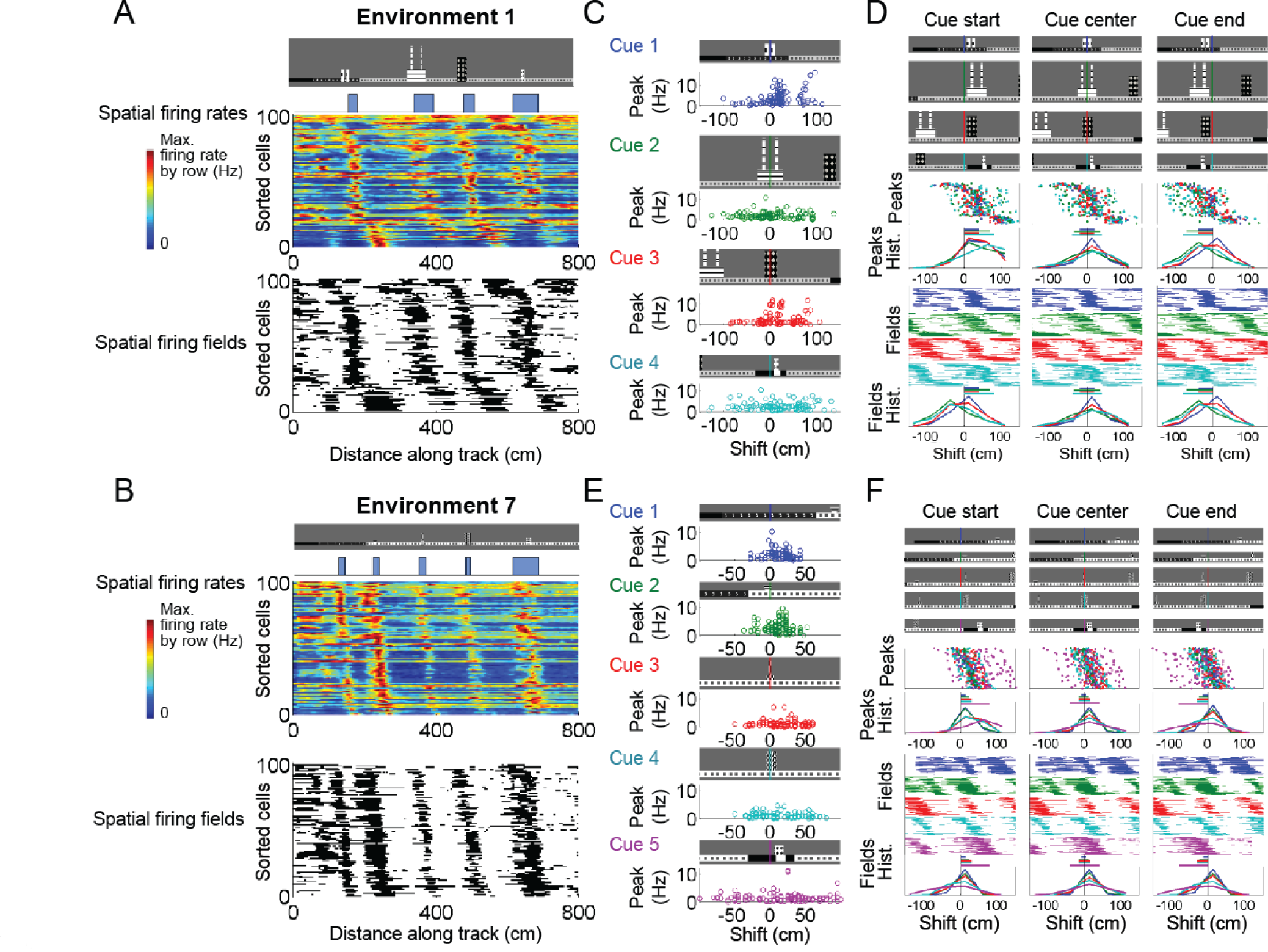
Cue cells form a sequence aligned at each cue. A. Cue cell sequence in an environment. Top two rows show a side view of the virtual track and the corresponding cue template below. The spatial firing rates (middle) and corresponding spatial firing fields (bottom) are shown for all cue cells recorded during navigation in a single environment. Each row is the firing rate of a single cell along the track, normalized by its maximum. The cells are sorted based on their spatial shifts calculated for alignment of spatial firing rates to the cue template (using the spatial shift of maximal correlation of the firing rate to the cue template, Materials and Methods). The firing fields are ordered in the same sequence as the firing rates. B. Sorted spatial firing rates and corresponding spatial firing fields for a different environment. C and E. The location of spatial firing rate peaks is plotted versus the peak heights for individual cues in the two environments in A and B, respectively. D and F. Alignment of cue cell sequences with different regions of cues in the environments in A and B, respectively. For each environment, the top panels are side views of sections of the environment around each cue (the earlier part of the track is to the left). Each row represents one cue. For each cue, the start, center, and end of the cue are centered in the left, middle, and right panels, respectively. Each cue is color-coded (vertical alignment bars in these plots and for the rest of the plots below, cue 1 = blue, cue 2 = green, cue 3 = red, cue 4 = cyan, cue 5 = purple). Since the cues were different sizes, the alignment of cue cell activity differed when aligned to the start, center, or end of the cues. For the alignments with different parts of each cue, the lower panels show the locations of the peaks of the cue cell spatial firing fields (Peaks), the distribution of the peaks (Peaks Hist.), the cue cell spatial fields (Fields), and the distribution of the spatial fields (Fields Hist). Both the spatial firing field peak and spatial field distributions show that the alignment to the center of the cues produced the most similar distributions of cue cell activity across all cues.

Since the virtual environment cues have various shapes and sizes, we next examined the alignment of cue cell spatial firing rates with each cue. We determined the location and amplitude of the peaks in the cue cell spatial firing rate and plotted these relative to each cue. We observed high peak amplitudes close to the cue locations for both tracks (Figures 4C and 4E). To determine how cue cell activity was aligned to cues, we also plotted cue cell spatial firing rate peaks and spatial firing fields aligned to the start, middle, and end of the cues to determine whether the sequences were more aligned to a particular region of the cue (start, middle, or end; Figures 4D and 4F). For each environment, we found the activity of all cue cells was best aligned to the center of the cue rather than the start or end of the cues (mean displacement ± standard error from position of alignment for the cue cell spatial firing rate peaks closest to each alignment location: cue start aligned: −49.0 cm ± 1.6, cue center aligned: 2.7 ± 1.3 cm, cue end aligned: 25.8 ± 1.1 cm; for cue cell spatial firing fields located within ± 50 cm of alignment location: cue start aligned: −9.0 ± 1.2 cm, cue center aligned: 2.2 ± 1.1 cm, cue end aligned: −6.5 ± 1.1 cm). Thus, cue cells, as a population, are activated in a sequence that is centered on prominent visual landmark locations.

### Cue cell pairwise activity patterns

Since cue cells are activated sequentially, do pairs of cue cells exhibit a correlation in their spike timing? We reasoned that temporal shifts (peaks in the spike time cross correlation function) should be observed between the firing of co-recorded cue cells that have peak activity at different spatial shifts in the sequence surrounding. Since the temporal correlation is a measurement of the spiking of these cells over time, this can be a useful measurement of the cue-independent activity of the population of cue cells, and thus represent an intrinsic relationship between cue cells in the MEC neural circuit.

We took all identified pairs of cue cells in our data and examined the spatial and temporal relationships of their spiking. Figure 5A shows several examples of the spiking of two cue cells recorded during ten runs on a virtual track. As expected from the cue cell spatial sequences described earlier, in many cases cue cell spatial firing patterns for pairs were offset relative to one another in space. To determine if there was a correlation of one cell to another in their spike timing, we calculated the spike time cross-correlogram for the spike times of each pair of cells. We observed that many pairs showed a temporal shift in the spike time cross-correlogram (Figure 5A, right). In Figure 5B, we calculated the relative spatial shift (peak in cross correlation of spatial firing rate) along with a temporal shift for each cue cell pair and plotted them in a 2D histogram. There is, as predicted, a general trend where the spatial and temporal shifts are correlated (Pearson correlation = 0.3).

**Figure 5.**
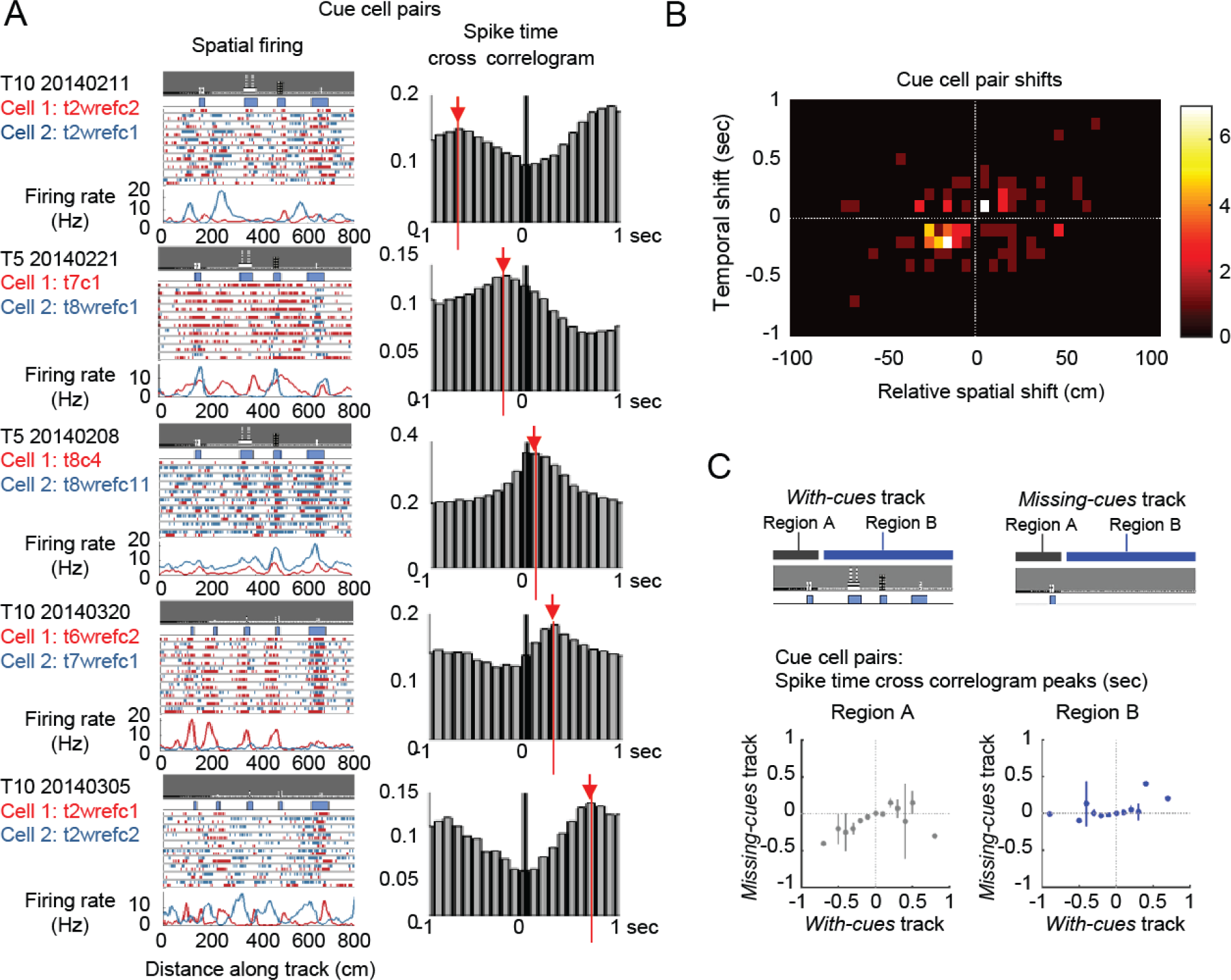
Cue cell pair activity and spatial and temporal shifts. A. Temporal activity relationship between pairs of cells with spatial offsets of their spatial firing rate. For each cell pair: left: the spiking activity for the two simultaneously recorded cue cells. The raster plot shows the spike locations on 10 runs along the track for each cell (cell 1 spikes are red, cell 2 spikes are blue). The spatial firing rates over all of the runs (not just for the ten runs shown) for both cells are plotted at the bottom. Right: the spike time cross-correlograms of the two cells with the temporal shift shown by a red line and arrow. B. Spatial and temporal shifts for all pairs of simultaneously recorded cue cells. The relative spatial shift was defined as the shift at which the correlation between the two firing rates had a local maximum closest to zero. The temporal shift was the shift at which the spike time cross-correlogram had a local maximum closest to zero. All cells are plotted in a histogram for relative spatial and temporal shifts, excluding zero values (no shift) along either axis. C. Temporal activity relationship between pairs of cells on *with-cues* and *missing-cues* tracks. Top: cartoons illustrating the track types and regions used for the data plotted below. For both tracks, the first region was identical in wall and cue locations (Region A). The second region (Region B) differed between the two tracks, one with cues (left, *with-cues* track) and one missing cues (right, *missing-cues* track). Data were sorted by different regions on the two tracks. Bottom: the mean +/− standard error of the temporal shifts between cue cell pairs recorded on both tracks. Bottom left: the time shifts of the peaks in the spike time cross-correlograms of cue cell pairs are plotted for the two tracks in Region A. Right: the time shifts for cue cell pairs on Region B of the two tracks. There was a range of time shift values when cues were present (for the ‘*with-cues* track’ plotted along the x-axis). However, there were fewer nonzero time shifts when cues were not present (see the distribution of values along the y-axis for the ‘m*issing-cues* track’ in Region B).

If the temporal correlations are produced by the sequential activation of the cells as cues are passed, then these correlations should disappear if the analysis is performed during times when no cues are present. Conversely, if correlations remain, this might provide an indication that the spike timing relationship between cells was produced independent of cues through intrinsic circuit mechanisms. Since we showed in Figure 5 that pairs of cue cells can be assigned both a spatial and temporal shift, we next asked whether there might still be a relationship between the spiking of the two cue cells even when cues are no longer present and the cells are active (despite the fact that no obvious sequence of activity is present in these regions without cues). Using a strategy similar to that used in Figure 2, we recorded from cue cells during virtual navigation on the *with-cues* and *missing-cues* tracks, and split the tracks into two regions: the start region where cues were identically present on both tracks (Region A), and the remaining region where cues were either present on *with-cues* track or missing on the *missing-cues* track (Region B). Using all cue cell pairs, we calculated the temporal shift for each pair in the four conditions (Regions A and B on both *with-cues* and *missing-cues* tracks). In Region A, there was a spread in the temporal shifts for pairs of cue cells and these shifts were correlated for the two tracks (Figure 5C left, Pearson correlation = 0.34). However, the temporal shifts in Region B of the two tracks were less correlated: while a similar spread of temporal shifts was observed when cue cells were recorded on the *with-cues* track (plotted along the x-axis of the bottom right panel in Figure 5C), but most cue cell pairs did not have a nonzero phase in the their relative spike timing when cues were missing (plotted along the y-axis of the bottom right panel in Figure 5C right, Pearson correlation = 0.13). This suggests that the spike timing relationship between cue cell pairs is present only when cues are present and thus when these cells are driven to be active in a sequential manner by locomotion past the cue.

### Side-preference of cue cells in superficial layers of the MEC

What type of sensory inputs could be driving the activity of cue cells? One potential candidate is visual input corresponding to spatial cues along the track, as suggested by the fact that the cue related spatial fields and the temporal shifts of cue cell spikes no longer existed when cues were removed from the track (Figures 2A and 5C). If visual input indeed drives cue cell responses, the cue cells in one hemisphere of the MEC, which receives inputs largely from the ipsilateral visual cortex (Olsen et al., 2017), should preferentially respond to cues on the contralateral side of the animal, as the visual cortex in one hemisphere generally receives visual information from the contralateral eye (Erskine and Herrera, 2014). To investigate this possibility, we designed an 18-meter long virtual track with asymmetric cues on the left and right sides of the track (different from the tracks used in above experiments, where identical visual cues were present on both sides of the track) and examined cue cell responses to these cues (Figure 6A). To increase the sampling and obtain precise anatomical information of cue cells in specific layers of the MEC, we used microprism-based cellular-resolution two-photon imaging to measure calcium responses of a large number of neurons in layers 2 and 3 of the MEC (Low et al., 2014). The genetically-encoded calcium indicator GCaMP6f was specifically expressed in layer 2 excitatory neurons of the MEC in the GP5.3 transgenic mice (Dana et al., 2014; Gu et al., 2018) and in layer 3 MEC neurons via viruses targeted to layer 3 of the MEC in wild type mice (Supplementary Figure 4). We only imaged neurons in the MEC on the left hemisphere.

**Figure 6.**
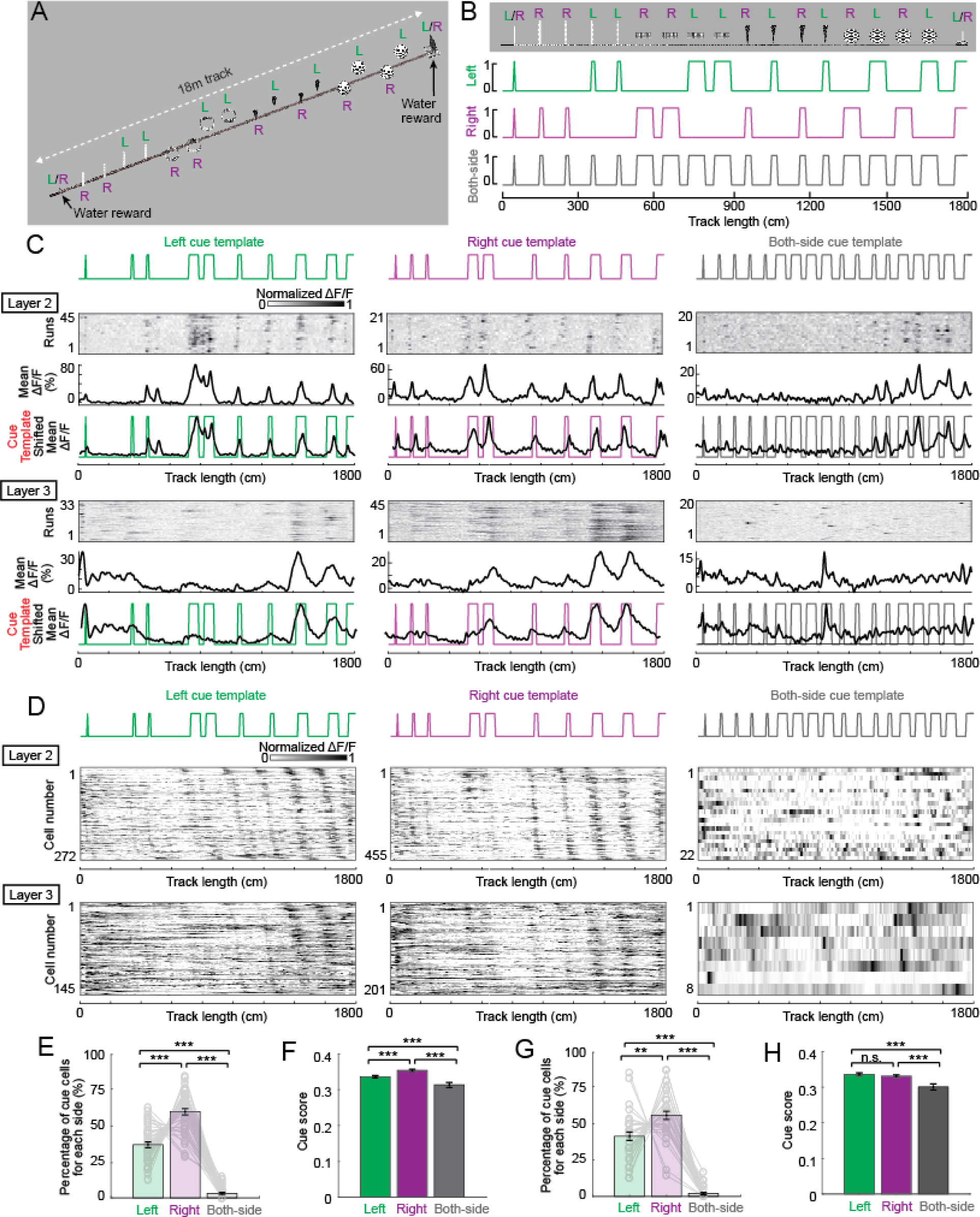
Cue cell responses to side-specific cues in superficial layers of the MEC. A. An 1800 cm (18 meter) long virtual track for imaging experiments. “L”, “R” and “L/R” indicate cues on the left, right and both sides of the track, respectively. B. Three types of cue templates corresponding to cues on the left, right, and both sides of the track. C. Examples of individual cue cells responding to the three side categories of cues in layers 2 and 3 of the MEC. For each cell: top: ΔF/F versus linear track position for a set of sequential traversals. Middle: mean ΔF/F versus linear track position. Bottom: overlay of the cue template and aligned mean ΔF/F (black) according to the shift, which gave the highest correlation between them (Materials and Methods). D. Cue cell sequences responding to the three side categories in layers 2 and 3. Each row is mean ΔF/F of a single cell along the track, normalized by its maximum. The cells are sorted by the spatial shifts of their mean ΔF/F to the cue template. E and F. Percentage (E) and cue scores (F) of cue cells in layer 2 responding to three side categories of cues. In E, each curve represents the percentage of left, right, and both-side cue cells (from left to right) in the whole cue cell population of a single FOV. Error bars: mean +/− SEM. All layer 2 cells shown in D were used in this analysis. G and H. Similar to E and F but for layer 3 cue cells. All layer 3 cells shown in C were used in this analysis.

During virtual navigation, we identified cue cells responding to the left, right, and both-side cues by calculating cue scores of their calcium responses along the virtual track with the three side-specific cue templates. Therefore, each cell had three cue scores. The highest score that passed the cue score threshold was used as the cue score of the cell, and the cell was thus assigned as a cue cell responding to the side of cues producing the highest score (Figure 6B). In both layers 2 and 3, we identified cells that responded to all three side categories of cues (Figure 6C) and formed sequences around these cues (Figure 6D). However, the majority of cue cells preferentially responded to cues on a single side (left or right) (96.9% and 98.0% for layers 2 and 3, respectively). While all imaging experiments were performed in the left hemisphere, there were significantly more right cue cells than left cue cells in both layers (Figure 6E and 6G). Also, in layer 2, the cue scores of the right cue cells were generally higher than those of the left cue cells, indicating the higher correlation of calcium responses of cue cells to right side cues (Figure 6F). This was not the case for layer 3 (Figure 6H). Overall, these results indicate that in superficial layers of the left MEC most cue cells respond to cues on the right side, strongly suggesting the role of side-specific visual input in driving the cue-related responses in these cells.

### Micro-organization of cue cells in the MEC

Cellular-resolution two-photon imaging provides detailed anatomical information of imaged neurons. This allowed for the study of the micro-organization of cue cells, providing additional information about potential connectivity between the cells and the inputs they receive. Since the majority of cue cells only responded to cues on single sides, we largely focused on right or left cue cells for this micro-organization study. A single field of view (FOV) in layer 2 or 3 generally contained multiple right and left cue cells (Figures 7A, B, D, and E). However, we observed that these cue cells tended to cluster, which was revealed by the shorter physical distances between cue cells than the distances between cue and non-cue cells (Figures 7C and 7F, left panels). In addition, cue cells with the same side preferences also tended to cluster, indicated by the fact that cue cells with the same side preferences generally showed shorter physical distances than those with different side preferences (Figures 7C and 7F, middle and right panels).

**Figure 7.**
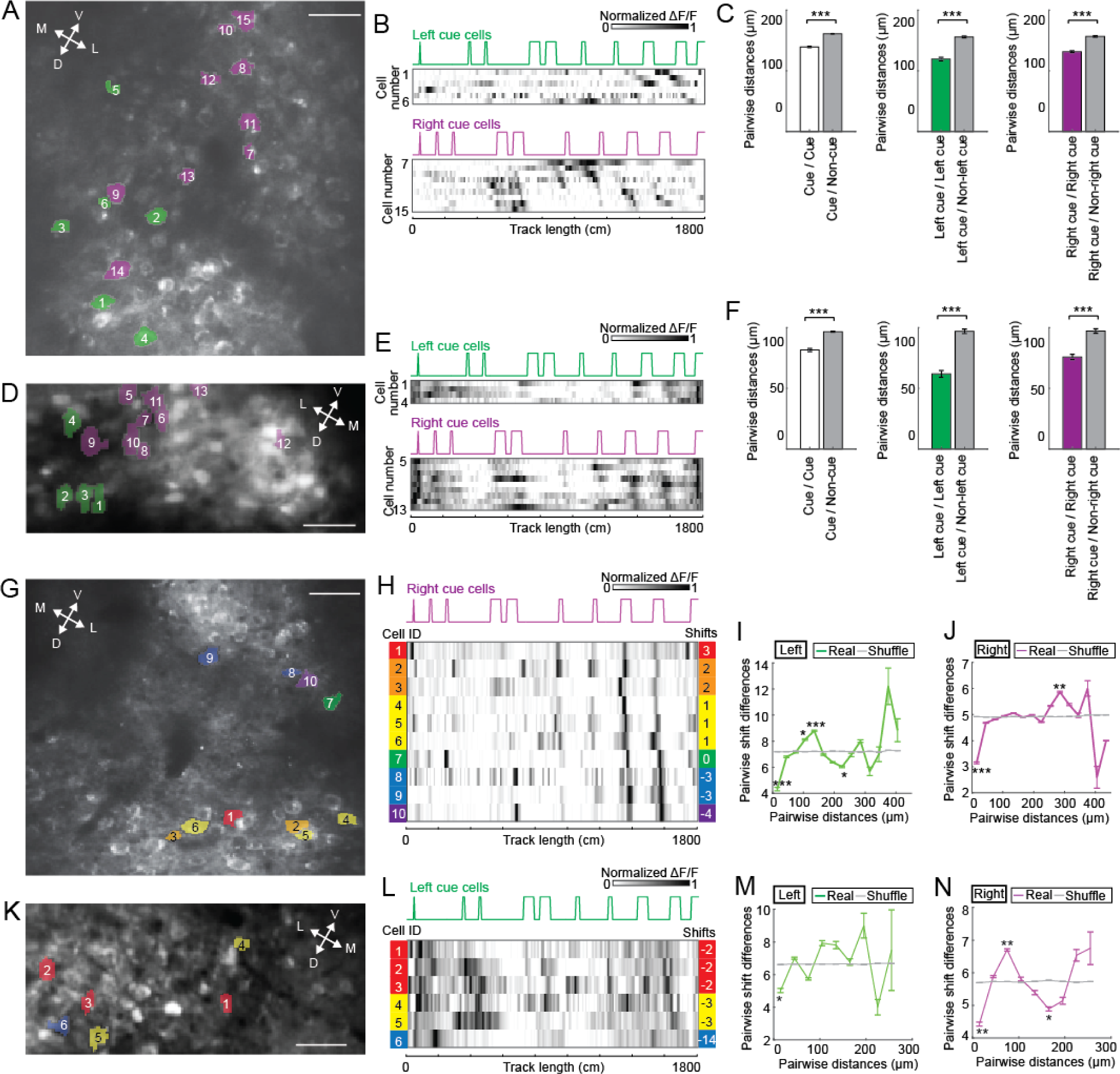
Micro-organization of cue cells in layers 2 and 3. A. A FOV in layer 2 of the MEC containing left (green) and right (magenta) cue cells. Cell indices are consistent with those in B. Anatomical orientations: D: dorsal; V: ventral; M: medial; L: lateral, same in the panels D, G, and K. B. Calcium responses of left and right cue cells shown in A. Each row is mean ΔF/F of a single cell along the track, normalized by its maximum. The cells are sorted by the shift of their mean ΔF/F to the cue template of their preferred side, same in panels E, H, and L. C. Pairwise distance between different cell types in layer 2. Non-cue cells included all cells in the MEC except cue cells. Non-left cue cells included all cue cells responding to right and both-side cues. Non-right cue cells included all cue cells responding to left and both-side cues. Error bars: mean +/− SEM. The analyses included 749 cue cells, 2170 non-cue cells, 272 left cue cells, 455 right cue cells and 22 both-side cue cells. D-F. Similar to A-C but for layer 3 cue cells. The analyses in F included 355 cue cells, 1569 non-cue cells, 145 left-cue cells, 201 right-cue cells and 9 both side cue cells. G. A FOV in layer 2 of the MEC containing right cue cells with different spatial shifts. Individual cue cells are color-coded according to their spatial shifts shown in H. Cell indices are consistent with H. H. Calcium responses of the right cue cells in G and their spatial shifts to cues on the right side of the track. I and J. Pairwise physical distance of left-(I) and right-cue cells (J) in layer 2 of the MEC versus pairwise difference of the spatial shifts of their mean ΔF/F to the left and right cue templates, respectively. Error bars: mean +/− SEM. 1026 left-cue cell pairs and 2940 right-cue cells were used. K and L. Similar to G and H but for left cue cells in a FOV in layer 3. M and N. Similar to I and J but for layer 3 cue cells. 351 left-cue cell pairs and 733 right-cue cells were used.

We also investigated the relationship between the physical distances of cue cells and the difference of the spatial shifts of their calcium responses relative to cue location. In layers 2 and 3, we consistently observed that anatomically adjacent cue cells (physical distances around 30 µm) showed more similar spatial shifts, whereas the relationship was more varied if cue cells were further apart (Figures 7G-7N). The similar cue responses of adjacent cue cells suggest that they may share similar inputs or be connected.

## Discussion

We have described a novel class of cells in MEC—termed cue cells—that were defined by a spatial firing patterns consisting of spatial firing fields located near prominent visual landmarks. When navigating a cue-rich virtual reality linear track, the population of cue cells formed a sequence of neural activity that was repeated at every landmark and aligned to the landmark centers. When cues were removed, these cells no longer exhibited a sequence of spatial firing fields but remained active, although typically at a lower maximum firing rate. There were spatial and temporal shifts between pairs of cue cells when cues were present, but these shifts disappeared in the absence of cues, consistent with the idea that their responses were driven primarily by sensory input rather than internal brain dynamics. The sensory input was likely visual input, as supported the fact that cue cells in the superficial layers of the left MEC mostly responded to cues on the right side of the track. In layers 2 and 3 of the MEC, all cue cells clustered, and more specifically, cue cells with the same side preferences also clustered. The spatial responses of anatomically adjacent cue cells had similar spatial shifts to cue patterns on their preferred side of the track. These properties of cue cells suggest that they could provide a source of spatial information in the local circuits of MEC that could be used in error correction in landmark rich environments.

### Cue cells and previously identified cell types

By recording during foraging in real arenas, we were able to determine how cue cells were related to previously identified MEC cell types (grid, head direction, border), all of which are defined by their activity in bounded 2D environments. We found that most cue cells were not grid or border cells, yet they did have noticeably stable, and somewhat irregular, spatial firing patterns in real arenas. Cue cells account for a significant fraction (~22 %) of the previously unexplained spatially-stable cells in MEC. While most cue cells were not grid or border cells, a vast majority of them had some orientation tuning (~50%). The high prevalence of head direction tuning for these cells suggests either that cue cells may receive inputs from traditional head direction cells, or that a head direction preference is present for cue cells because of the location of particular features of the real arena that drives the activation of cue cells. Further work is required to determine the circuit mechanisms of this orientation tuning preference.

Do cue cells resemble spatially modulated cells in other brain regions, such as place cells or boundary vector cells? Place cells typically have only a single firing field during navigation along linear tracks, even in virtual reality environments with prominent visual cues along the tracks (Dombeck et al., 2010). This is distinctly different from cue cells, in which the number of spatial firing fields scales with the number of cues. Boundary vector cells, which were found in subiculum, encode distance to a boundary (Lever et al., 2009; Stewart et al., 2014). An identified boundary vector cell must have a spatial firing field that is uniformly displaced from a particular region of the boundary. The width of the spatial firing field is proportional to the distance from the boundary, meaning that cells shifted significantly from the border would have very wide spatial firing fields, which could cover a large majority of the environment (Lever et al., 2009; Stewart et al., 2014). Border cells are a special case of boundary vector cells. To determine whether cue cells might be boundary vector cells, we sorted their spatial responses in the real 2D arena based on the shifts of their spatial firing rates from the visual cue pattern on a virtual track (Supplementary Figure 2). We found no obvious trends in the spatial firing patterns of these cue cells in the real arena. Along with this, many cells had multiple fields or fields that were not uniformly displaced from the border of the environment. These spatial firing field features of cue cells were inconsistent with those of boundary vector cells. Thus, cue cells have properties distinct from both place cells and boundary vector cells. Their spatial firing responses were distributed throughout the arena, despite there being a single white cue card visible, indicating that these cells perform more complex computations in real arenas where spatial cues take many forms in comparison to the cues along a virtual track. Previous studies have also found similar complex responses of non-grid cells in the MEC encoding features of real environments (Diehl et al., 2017; Hardcastle et al., 2017).

### Cue cells and path integration

It has been hypothesized that the MEC is a central component of a path integrator that uses self-motion information to update a spatial metric encoded by the population of grid cells (Burak and Fiete, 2009; Fuhs and Touretzky, 2006; McNaughton et al., 2006). Grid cells are grouped into modules based on each cell’s grid spacing. Each grid module maintains its own orientation, to which all grid cells align and are related by a two-dimensional spatial phase. Grid cells in a given module maintain their relative spatial phase offsets across different environments (Fyhn et al., 2007), including linear tracks (Yoon et al., 2016), indicating that the population of grid cells form a consistent spatial metric largely defined only by the two-dimensional phase. This provides support for the idea that grid cell dynamics are constrained to a two-dimensional attractor manifold (Yoon et al., 2013). In a manifold-based path integrator, spatial location is represented as the location of the grid cell population activity on the attractor manifold. Self-motion signals, such as running speed in a particular direction, move the population activity along the manifold, such that changes in location are proportional to the integral of the velocity over time. Path integration is inherently a noisy process that requires calibration and error correction for more accurate estimates of position.

In the context of continuous attractor models for path integration, it is interesting to consider the potential functional roles of cue cells. One role could be to act as external error-correction inputs to the path integrator network that tend to drive the neural activity pattern to manifold locations appropriate for each landmark. An analogous use was proposed for border cells, in which they contribute to error correction near boundaries (Hardcastle et al., 2015; Pollock, 2018). In this role, it would be advantageous if the cue cell population activity represented distinct cues differently, so that more refined information about the locations of individual cues might be coded in cue cell firing and used in the correction. In Supplementary Figure 5, we provide preliminary evidence that the population activity of cue cells encoded information about the particular identity of each cue. More work is needed to further characterize the precise nature of this precise coding of unique landmarks and, with new models, to determine how effectively it might be used to drive an attractor network to the appropriate spatial locations when interacting with a noisy path integrator.

An alternative, or additional, role for cue cells in path integration would be to produce a continuous adjustment of location. The sequence of activity that was produced across the cue cell population as individual landmarks were passed could drive the network activity continuously along the manifold, in essence acting as a velocity input that is quite different from those traditionally considered, such as running speed. This use, as an effective velocity, is analogous to the recent demonstration (Hopfield, 2015; Ocko, 2018) that the collective state of a line attractor can be moved continuously along the manifold by an appropriately learned sequence of external inputs. In essence, the set of inputs at each time point move the location of the activity on the attractor a slight amount, and this is repeated continuously to produce smooth motion, without requiring asymmetric synaptic connectivity and velocity-encoding signals of previous path integrator models (Burak and Fiete, 2009; Fuhs and Touretzky, 2006; McNaughton et al., 2006; Ocko, 2018). In principle, both path-integration and error correction can be combined through this process.

### Cue cells in real and virtual environments

Why cue cells had such stable and easily classified activity patterns in virtual reality, but not in the real arena remains an open question. Navigating along a virtual track differs greatly from foraging in a complex real arena. In the case of virtual reality, the animal encounters a single cue at a time, so the representation of location using visual information is primarily from a limited orientation to individual cues, one pair at a time, along the track. It is possible that forming a representation of location in the real arena requires triangulation from many cues located at various angles and distances away. Despite the simple design of our real arena with a single cue card on one wall, there could be multimodal features from the floor, walls, or from distal cues outside of the arena. Navigation along simple virtual environments comprised of only visual cues sets up an ideal experimental paradigm to further understand the activity of these cells. Future experiments could probe other features of the cells with more perturbations of the virtual environment.

### Cell classes in MEC

Although our analysis and discussion of cue cells have largely followed the traditional approach of describing MEC cells according to discrete classes, it is interesting to note that cue scores, like grid, head direction, and border scores, each form a continuum and that a significant fraction of cells in MEC are conjunctive for more than one class (Figure 3). The conjunctive coding in neural firing in MEC is also demonstrated by a recent study (Hardcastle et al., 2017) and is conceptually analogous to the “mixed selectivity” in neural codes that have been increasingly recognized in cognitive, sensory and motor systems (Finkelstein et al., 2015; Fusi et al., 2016; Rigotti et al., 2013; Rubin et al., 2014). Recently, mixed selectivity has been demonstrated to be computationally useful in evidence integration and decision-making by allowing the selection of specific integrating modes in accumulating evidence to guide future behavior (Mante et al., 2013; Ulanovsky and Moss, 2011). Reframing this in the context of path integration, it will be useful to determine how navigation systems might use mixed selectivity and context-specific integrating modes to weigh different accumulating information (different velocity and position inputs) according to the reliability of that information during navigation in complex, feature-rich environments.

## Materials and Methods

### Animals

All procedures were approved by the Princeton University Institutional Animal Care and Use Committee and were in compliance with the Guide for the Care and Use of Laboratory Animals. Four C57BL/6J male mice, 3-6 months old, were used for electrophysiological experiments. Two 10-week old male mice were used for the two-photon imaging of layer 3 neurons. Mice used for the two-photon imaging of layer 2 neurons were six 10-to 12-week old GP5.3 males, which were heterozygous for carrying the transgene Thy1-GCaMP6f-WPRE to drive the expression of GCaMP6f (Dana et al., 2014).

### Experimental Design and Statistical Analysis

All electrophysiological data are represented as mean ± STD and imaging data are represented as mean ± SEM, unless otherwise noted. A student’s t-test was always used to evaluate whether the difference of two groups of values was statistically significant. Significance was defined using a p value threshold of 0.05 (* p<0.05, ** p<0.01, *** p<0.001). All analysis was performed using custom Matlab software and built in toolkits. All correlations were Pearson correlations unless otherwise specified.

### Code Accessibility

Code for all the analyses will be available upon request.

### Real arena for tetrode recording

Experiments were performed as described previously (Domnisoru et al., 2013). The real arena consisted of a 0.5 m × 0.5 m square enclosure with black walls at least 30 cm high and a single white cue card on one wall. Animals foraged for small pieces of chocolate (Hershey’s milk chocolate) scattered throughout the arena at random times. Trials lasted 10-20 minutes. On each recording day, real arena experiments were always performed before virtual reality experiments. Video tracking was performed as described previously (Domnisoru et al., 2013) using a Neuralynx acquisition system (Digital Lynx). Digital timing signals, which were sent and acquired using NI-DAQ cards, and controlled using ViRMEn software in Matlab (Aronov and Tank, 2014) were used to synchronize all computers.

### Virtual reality (VR)

The virtual reality system was similar to those described previously (Dombeck et al., 2010; Domnisoru et al., 2013; Gauthier and Tank, 2018; Gu et al., 2018; Harvey et al., 2012; Harvey et al., 2009; Low et al., 2014). ViRMEn software (Aronov and Tank, 2014) was used to design the linear VR environment, control the projection of the virtual world onto the toroidal screen, deliver water rewards (4µl) through the control of a solenoid valve, and monitor running velocity of the mice. Upon running to the end of the track, mice were teleported back to the beginning of the track.

#### VR for tetrode recording

The animal ran on a cylindrical treadmill, and the rotational velocity of the treadmill, which was proportional to mouse velocity, was measured using sequential sampling of an angular encoder (US Digital) on each ViRMEn iteration (~60 iterations per second). The tracks were 8 meters long with identical cues on both side of the track.

#### VR for imaging

Mice ran on an air-supported spherical treadmill, which only rotated in the forward/backward direction. Their heads were held fixed under a two-photon microscope (Gu et al., 2018; Low et al., 2014). The motion of the ball was measured using an optical motion sensor (ADNS3080; red LED illumination) controlled with an Arduino Due. The VR environment was rendered in blue and projected through a blue filter (Edmund Optics 54-462). The track was 18 meters long with asymmetric cues on two sides of the track. Water rewards (4 µl) were delivered at the beginning and the end of the track.

### Microdrives and electrode recording system

Custom microdrives and the electrophysiology recording system used were similar to those described previously (Aronov and Tank, 2014; Domnisoru et al., 2013; Kloosterman et al., 2009). Tetrodes were made of PtIr (18 micron, California Fine Wire) and plated using Platinum Black (Neuralynx) to 100-150 kΩ at 1 kHz. A reference wire (0.004” coated PtIr, 0.002” uncoated 300 µm top) was inserted into the brain medial to the MEC on each side, and a ground screw or wire was implanted near the midline over the cerebellum.

The headstage design was identical to the one used previously (Aronov and Tank, 2014) with the addition of solder pads to power two LEDs for use in tracking animal location and head orientation. Custom electrode interface boards (EIBs) were also designed to fit within miniature custom microdrives. A lightweight 9-wire cable (Omnetics) connected the headstage to an interface board. The cable was long enough (~3m) to accommodate the moving of the animal between the real arena and the virtual reality system without disconnection.

### Surgery

#### Tetrode recording

Surgery was performed using aseptic techniques, similar to those described previously (Domnisoru et al., 2013). The headplate and microdrive were implanted in a single surgery that lasted no longer than 3 hours. Bilateral craniotomies were performed with a dental drill at 3.2 mm lateral of the midline and just rostral to the lambdoid suture. After the microdrive implantation, 4-6 turns were slowly made on each drive screw, lowering the tetrodes at least 1 mm into the brain. Animals woke up within ~10 minutes after the anesthesia was removed and were then able to move around and lift their heads.

#### Imaging

The surgical procedures were similar to those described previously (Low et al., 2014). A microprism implant was composed of a right angle microprism (1.5 mm side length, BK7 glass, hypotenuse coated with aluminum; Optosigma), a circular coverslip (3.0 mm diameter, #1 thickness, BK7 glass; Warner Instruments) and a thin metal cylinder (304 stainless steel, 0.8 mm height, 3.0 mm outer diameter, 2.8 mm inner diameter; MicroGroup) bonded together using UV-curing optical adhesive (Norland #81). The microprism implantation was always performed in the left hemisphere (Gu et al., 2018; Low et al., 2014). A circular craniotomy (3 mm diameter) was centered 3.4 mm lateral to the midline and 0.75 mm posterior to the center of the transverse sinus (at 3.4 mm lateral). The dura over the cerebellum was removed. The microprism assembly was manually implanted, with the prism inserted into the subdural space within the transverse fissure. The implant was bonded to the skull using Vetbond (3M) and Metabond (Parkell). A titanium headplate with a single flange was bonded to the skull on the side opposite to the side of the craniotomy using Metabond. For imaging layer 3 neurons in the MEC, AAV1.hSyn.GCaMP6f.WPRE.SV40 (Penn Vector Core) virus was diluted 1:4 in a solution of 20% (w/v) mannitol in PBS and pressure injected at two sites (200 nl/site): (1) ML 3.00 mm, AP 0.77 mm, depth 1.79 mm; (2) ML 3.36 mm, AP 0.60 mm, depth 1.42 mm.

### Two-photon imaging during virtual navigation

Imaging was performed using a custom-built, VR-compatible two-photon microscope (Low et al., 2014) with a rotatable objective. The 920 nm excitation laser was delivered by a mode-locked Ti:sapphire laser (Chameleon Ultra II, Coherent, 140fs pulses at 80 MHz). The laser scanning for imaging layer 2 neurons of the MEC was achieved by a resonant scanning mirror (Cambridge Tech.). The laser scanning for imaging layer 3 neurons of the MEC was achieved by a galvanometer XY scanner (Cambridge Tech.). Fluorescence of GCaMP6f was isolated using a bandpass emission filter (542/50 nm, Semrock) and detected using GaAsP photomultiplier tubes (1077PA–40, Hamamatsu). The two objectives used for imaging layers 2 and 3 were Olympus 40×, 0.8 NA (water) and Olympus LUCPLFLN 40x, 0.6 NA (air), respectively. Ultrasound transmission gel (Sonigel, refractive index: 1.3359 (Larson et al., 2011); Mettler Electronics) was used as the immersion medium for the water immersion objective used for layer 2 imaging. The optical axes of the microscope objective and microprism were aligned at the beginning of each experiment as described previously (Low et al., 2014). Microscope control and image acquisition were performed using ScanImage software (layer 2 imaging: v5; layer 3 imaging: v3.8; Vidrio Technologies (Pologruto et al., 2003)). Images were acquired at 30 Hz at a resolution of 512 x 512 pixels (~410 x 410 µm FOV) for layer 2 imaging, and 13 Hz at a resolution of 64 x 256 pixels (~100 x 360 µm FOV) for layer 3 imaging. Imaging and behavioral data were synchronized by simultaneously recording the voltage command signal to the galvanometer together with behavioral data from the VR system at a sampling rate of 1 kHz, using a Digidata/Clampex acquisition system (Molecular Devices).

### Histology

#### For tetrode recording

To identify tetrode locations, small lesions were made by passing anodal current (15 µA, 1 sec) through one wire on each tetrode. Animals were then given an overdose of Ketamine (200 mg/kg)/Xylazine (20 mg/kg) and perfused transcardially with 4% formaldehyde in 1X PBS. At the end of perfusion, the microdrive/headplate assembly was carefully detached from the animal. The brain was harvested and placed in 4% formaldehyde in 1X PBS for a day and then transferred to 1X PBS. To locate tetrode tracks and lesion sites, the brain was embedded in 4% agarose and sliced in 80 µm thick sagittal sections. Slices were stained with a fluorescent Nissl stain (NeuroTrace, Molecular Probes), and images were acquired on an epifluorescence microscope (Leica) and later compared with the mouse brain atlas (Paxinos). To identify which tetrode track belonged to each tetrode of the microdrive, the microdrive/headplate assembly was observed with a microscope to determine the location of each tetrode in the cannula, the relative lengths of the tetrodes, and whether the tetrodes were parallel or twisted. If tetrodes were twisted, then recordings were used only if grid cells were found on the tetrode.

#### For imaging

For verifying the layer-specific expression of GCaMP6f in the MEC, animals were transcardially perfused, as described above, and their brains were sliced in 100 µm thick sagittal sections. A fluorescent Nissl stain was performed as described above.

### General data processing for tetrode recording

Data analysis was performed offline using custom Matlab code. Electrophysiology data were first demultiplexed and filtered (500 Hz highpass). Spikes were then detected using a negative threshold set to three times the standard deviation of the signal averaged across electrodes on the same tetrodes. Waveforms were extracted and features were then calculated. These features included the baseline-to-peak amplitudes of the waveforms on each of the tetrode wires as well as the top three principal components calculated from a concatenation of the waveforms from all wires.

#### Cluster separation

Features of the waveforms were plotted with a custom Matlab GUI. Criteria for eliminating clusters from the dataset were: units with less than 100 spikes (in real arena or virtual tracks), the minimum spatial firing rate along the virtual track > 10 Hz or the maximum firing rate > 50 Hz. After this, 2825 clusters remained. Since clusters were cut with two different methods (using all 4 electrodes and using 3 electrodes with the fourth subtracted as a reference), repeats needed to be removed from the overall dataset. Repeats were found using a combination of 3 measures: the Pearson correlation of the real arena spatial firing rate, the same correlation of the virtual track spatial firing rate and the ISI distribution of the spikes merged between the two clusters. If the sum of these scores exceeded 2.25 then the clusters were considered to be from the same cell; the cluster with the larger number of spikes was kept, and the other cluster was discarded.

Recordings were performed on four animals over two months. From these recordings, 5940 clusters were manually cut using a custom Matlab GUI. Of these, there were 1081 clusters that were identified on tetrodes that were histologically identified to be in MEC during the recording that also passed our cluster quality criteria. The grid scores of these cells were calculated and a grid score threshold was calculated using shuffled permutations of these cells (Aronov and Tank, 2014; Domnisoru et al., 2013). Any tetrode on a particular day with a grid cell was then added to the database from that date on. The final database contained 1590 clusters.

### Spatial firing rates of tetrode data

Position data (including head orientation in real 2D arenas) were subsampled at 50 Hz and spikes were assigned into the corresponding 0.02 sec bins. Velocity was calculated by smoothing the instantaneous velocity with a moving boxcar window of 1 second. Only data in which the animal’s smoothed velocity exceeded 1 cm/sec were used for further analyses of firing rates or scores.

#### Real arena spatial firing rate

2D arenas were divided into 2.5x2.5 cm bins. Spike counts and the total amount of times spent in these bins were convolved with a Gaussian window (5x5 bins, σ = 1 bin). Firing rate was not defined for spatial bins visited for a total of less than 0.3 seconds.

#### Real arena, head direction

The animal’s head direction was binned in 3-degree intervals. For each angle bin, the spike count and the total amount of time spent (occupancy) was calculated. These values were separately smoothed with a 15 degree (5 bins) boxcar window, and the firing rate was computed as the ratio of the smoothed spike count to the smoothed occupancy.

#### Virtual track spatial firing rate

Virtual tracks were divided into 5 cm bins. Spike counts and the amount of time spent in these bins were smoothed independently with a Gaussian window (3 point, σ = 1). The smoothed firing rate was calculated as the smoothed spike position distribution divided by the smoothed overall position distribution.

#### Spatial firing fields

To calculate spatial firing fields we created time arrays for position and for number of spikes of a cell. Time bins were 100 msec. For position, we calculated the average position within each chunk of time (5 data points since data were interpolated to 20 msec sampling intervals). We then divided the spatial track into 5 cm bins and determined in which bin the average position was located. For spikes, we counted the number of spikes in that 100 msec interval. This generated two arrays in time (sampled at 100 msec), one with spike count and one with spatial bin location along the track. We then circularly permuted the spike count array by a random time interval between 0.05 x recording length and 0.95 x recording length. We then calculated the smoothed firing rate of this shuffled spike time array with the spatial bin location array. This was repeated 100 times, and the shuffled spatial firing rate was calculated for each permutation. The p-value was defined for each spatial bin along the track as the fraction of permutations on which the firing rate in that bin was above the actual firing rate. Any bin in which the p-value was less than 0.3 was considered part of a firing field.

#### Firing field distributions

For each cue cell, we define an array (5 cm bins) that is 1 when there is a firing field and 0 otherwise. To look at the distribution of firing fields for the population of cells, we sum the values for each bin across all cue cells and divide by the number of cells. This gives the fraction of cue cells with firing fields at each location. The plot of this fraction versus location was defined as the population firing field distribution.

### Scores for cells in tetrode data

For all scores below, 400 shuffles were performed with spike times circularly permuted by a random amount of time chosen between 0.5 x recording length and 0.95 x recording length, a standard method for determining score thresholds (Domnisoru et al., 2013). Shuffled distributions from all units combined were used to calculate a threshold at 95th percentile.

#### Grid score

The unbiased autocorrelation of the 2D firing rate in a real arena was first calculated (Hafting et al., 2005). Starting from the center of the 2D autocorrelation function, an inner radius was defined as the smallest radius of three different values: local minimum of the radial autocorrelation; where the autocorrelation was negative; or at 10 cm. Multiple outer radii were used from the inner radius + 4 bins to the size of the autocorrelation - 4 bins in steps of 1 bin. For each of these outer radii, an annulus was defined between the inner and the outer annulus. We then computed the Pearson correlation between each of these annuli and its rotation in 30 degree intervals from 30 to 150 degrees. For each annulus we then calculated the difference between the maximum of all 60 and 120 rotation correlations and the minimum of all 30, 90, and 150 degree correlations. The grid score was defined to be the maximum of all of these values across all annuli.

#### Head direction score

The head direction score was defined to be the mean vector length of the head direction firing rate (Giocomo et al., 2014). The head direction angle was defined to be the orientation of the mean vector of the head direction firing rate.

#### Spatial/head direction stability

This was calculated as described previously (Boccara et al., 2010). Recording sessions were divided into two parts, the firing rate was calculated for each, and the spatial stability was defined as the Pearson correlation between the two parts.

#### Cue score

The cue score was developed to measure the correlation of the spatial firing rate to the visual cues of the environment. A “cue template” was defined in 5 cm bins with value equal to 1 for bins that included the area between the front and back edges of each cue and 0 elsewhere. The cross correlation between the cue template and the firing rate was first calculated (relative shift <= 300 cm). The peak in the cross correlation with the smallest absolute shift from zero was chosen as the best correlation of the firing rate to the cue template. The spatial shift at which this peak occurred was then used to displace the cue template to best align with the firing rate. The correlation was then calculated locally for each cue. The local window included the cue and regions on either side extending by half of the cue width. The mean of local correlation values across all cues was calculated and defined as the “cue score”. An illustration of this method is shown in Figure 1B. This score effectively distinguished grid cells from cue cells, because grid cells generally did not have peaks at consistent locations relative to all the cues. The small number of grid cells that passed the cue score shuffle test also tended to have activity in other locations, where cues were not present.

#### Ridge/background ratio

The ridge/background ratio was calculated on the smoothed spatial firing rate at each cue location. The spatial firing rate of each cell was shifted to maximally align to the cue template as was done to calculate the cue score. The 5 bins (25 cm) in the center of each cue location are defined to be bins for the ridge. Background bins were all bins outside of cue locations displaced in both directions by [cue half-width + 20] to [cue half-width + 30]. For each cue, the ridge/background ratio was calculated as the mean firing rate in the ridge bins divided by the mean firing rate in the background bins. The ridge/background ratio for the cell was defined to be the mean of these individual ridge/background ratios. We performed 1000 shuffles of the data, as described above, and calculated the mean ridge/background ratio for each shuffle. The p value is the (number of shuffled data mean values larger than the mean ridge/background ratio of the data)/(number of shuffled data mean values less than the mean ridge/background ratio of the data).

### General imaging data processing

All imaging data were motion corrected using a whole-frame, cross-correlation-based method (Dombeck et al., 2010) and were then used to identify regions of interest (ROIs) using an independent component analysis (ICA) based algorithm (Mukamel et al., 2009) (for individual layer 3 field of view (FOV): mu = 1, 150 principal components, 150 independent components, s.d. threshold = 3; for individual layer 2 FOV, which was evenly split as nine blocks before ICA: mu = 0.7, 30 principal components, 150 independent components, s.d. threshold = 3). Fluorescence time series of these ROIs were extracted from all motion-corrected stacks. The fractional change in fluorescence with respect to baseline (ΔF/F) was calculated as (F(t) – F_0_(t)) / F_0_(t), similar to what was described previously (Gu et al., 2018; Low et al., 2014). Significant calcium transients were identified as those that exceeded cell-specific amplitude/duration thresholds (so that artefactual fluctuations were expected to account for less than 1% of detected transients (Dombeck et al., 2007)). Mean ΔF/F of the whole imaging session or individual traversals was calculated as a function of position along the virtual track for non-overlapping 5 cm bins. Only data points during which the mouse’s running speed met or exceeded 1 cm/s were used for the calculation.

### Identifying cue cells in imaging data

#### Selection of cells

candidates for cue cells were restricted to cells that contained at least one in-field period and one out-of-field period based on a p-value analysis of their calcium responses (Domnisoru et al., 2013; Gu et al., 2018; Heys et al., 2014; Yoon et al., 2016). Similar to identifying spatial fields for tetrode-recorded cells, in‐ and out-of-field periods were defined by comparing the mean ΔF/F value in each 5 cm bin to that of a random distribution created by 1000 bootstrapped shuffled responses, which were generated by rotating the ΔF/F trace starting from random sample numbers between 0.05 × N_samples_ and 0.95 × N_samples_ (N_samples_: number of samples in the ΔF/F trace). For each bin, the p-value equaled the percent of shuffled mean ΔF/F that were above the real mean ΔF/F. In-field-periods were defined as three or more adjacent bins (except at the beginning and end of the track where two adjacent bins were sufficient) whose p-value ≤ 0.2 and for which at least 10% of the runs contained significant calcium transients within the period. Out-of-field periods were defined as two or more bins whose p-value ≥ 0.75.

#### Defining cue cells responding to left, right, and both-side cues

Three types of cue template were generated: left, right, and both side cue templates corresponded to cues localized on the left, right, and both sides of the track, respectively. Three cue scores for each cell were calculated as described above according to the three types of template. The highest score and its corresponding side were used as the cue score and side preference for that cell, respectively. The cue score of a cell was further compared to those of bootstrapped shuffled responses (200 shuffles for each cell, generated as described above). Cue scores of all shuffles from all cells to all types of cue templates were combined together. If the cue score of a cell exceeded the 95^th^ percentile of the shuffled scores, the cell was a cue cell responding to cues with that side preference.

## Acknowledgements

We thank current and former members of the Tank lab, Ila Fiete, and Anika Kinkhabwala for helpful discussions, and Jeffrey Santner and Alexander Riordan for comments on the manuscript. This work was supported by NINDS Grant 5R37NS081242 (D.W.T), NIMH Grant 5R01MH083686 (D.W.T), NIH Postdoctoral Fellowship Grant F32NS070514-01A1 (A.A.K).

## Figures and figure legends

**Supplementary Figure 1.**
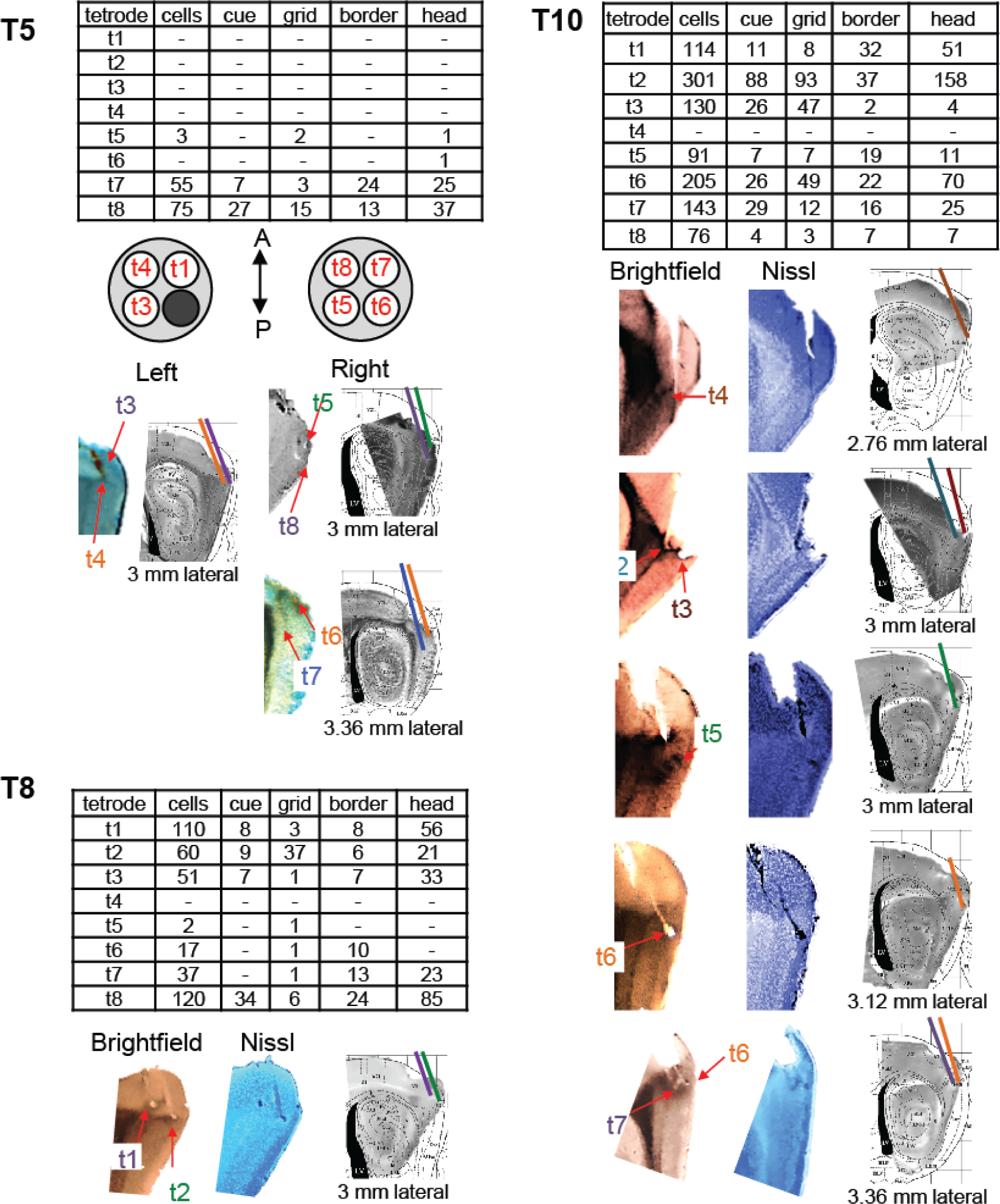
Histology. For each animal, T5, T8, and T10, a summary table shows the number of cells in the database from each tetrode, as well as the number of cue, grid, border and head direction cells within that population. For T5, an example of the orientation of the tetrodes within the cannula on either side of the brain is shown. This was used as a guide to identify individual tetrodes based on the tetrode tracks and lesions in the brain. The images of sagittal brain sections taken under the bright field microscope and the fluorescence microscope after fluorescent Nissl stain are shown for individual animals. Individual tetrodes were identified, color-coded and labeled in the figure. All images were sharpened and recolored to emphasize tetrode tracks and lesions.

**Supplementary Figure 2.**
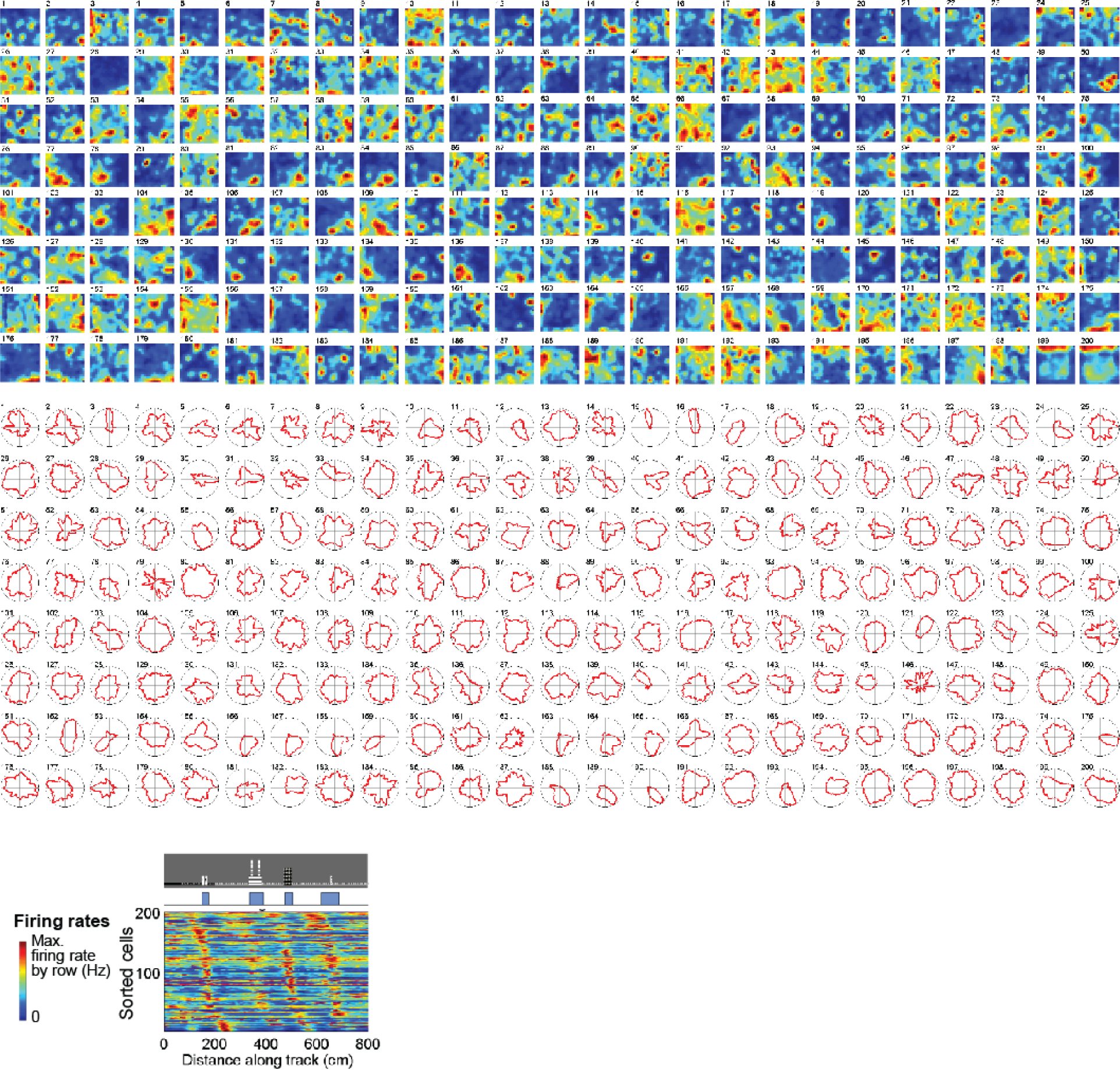
Cue cell activity in real arenas. Top and middle panels: the spatial and head direction firing rates of cue cells in a real arena are sorted based on the spatial shifts of their spatial firing fields to the cue template in virtual reality (bottom panel). No clear patterns of changes in number, size and location of firing fields or the mean vector length of head direction firing rates were observed.

**Supplementary Figure 3.**
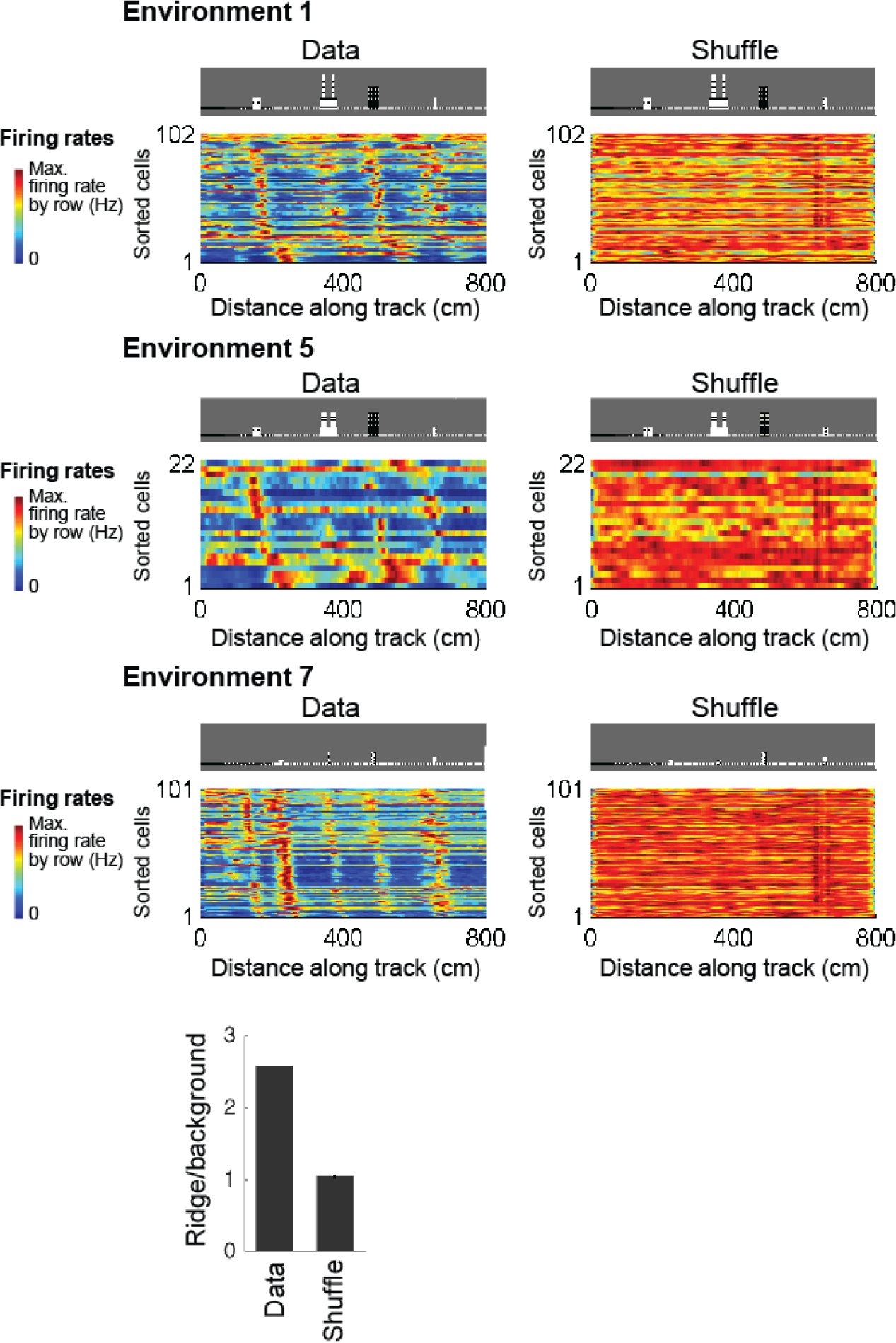
Cue cell sequences and the ridge/background ratio compared to shuffled data. For each environment, left: spatial firing rates (normalized by each cell’s maximum firing rate) of cue cells were sorted by their shifts relative to the cue template; right: an example of one set of shuffled firing rates of all cue cells. The shuffled spatial firing rates were also sorted by the spatial shifts relative to the cue template. Bottom: mean ridge/background ratios for all cue cells (2.58, left) and all shuffles (1.04+/− 0.003, right). Error bars: mean +/− STD.

**Supplementary Figure 4.**
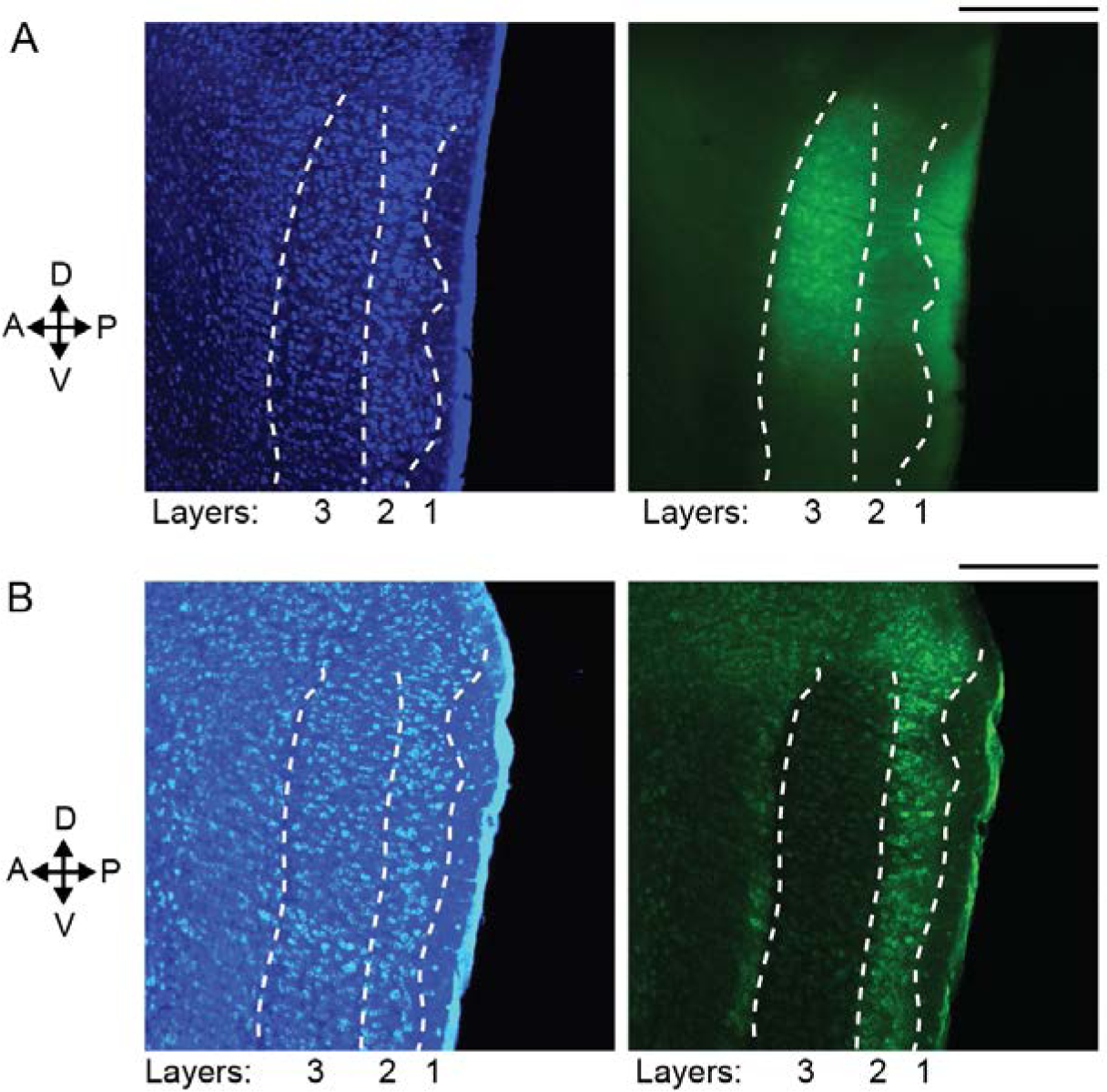
Layer-specific expression of GCaMP6f in the mouse MEC. A. Expression of GCaMP6f in layer 3 of the MEC, shown by a sagittal brain slice of a wild type mouse, prepared 13 days after the injection of AAV2/1-hSyn-GCaMP6f virus to the MEC. Left: blue epifluorescence image showing cell bodies in layers 2 and 3 (between white dotted curves) of the MEC labeled by fluorescent Nissl staining. Right: green epifluorescence image of the same slice on the left showing GCaMP6f expression of dorsal layer 3 neurons in the MEC. Scale bar: 200 µm. B. Expression of GCaMP6f in layer 2 of the MEC, shown by a sagittal brain slice of a GP5.3 mouse, 5 months of age. Left: blue epifluorescence image showing cell bodies in layers 2 and 3 (between white dotted curves) of the MEC labeled by fluorescent Nissl staining. Right: green epifluorescence image of the same slice on the left showing GCaMP6f expression of layer 2 neurons in the MEC. Scale bar: 200 µm.

**Supplementary Figure 5.**
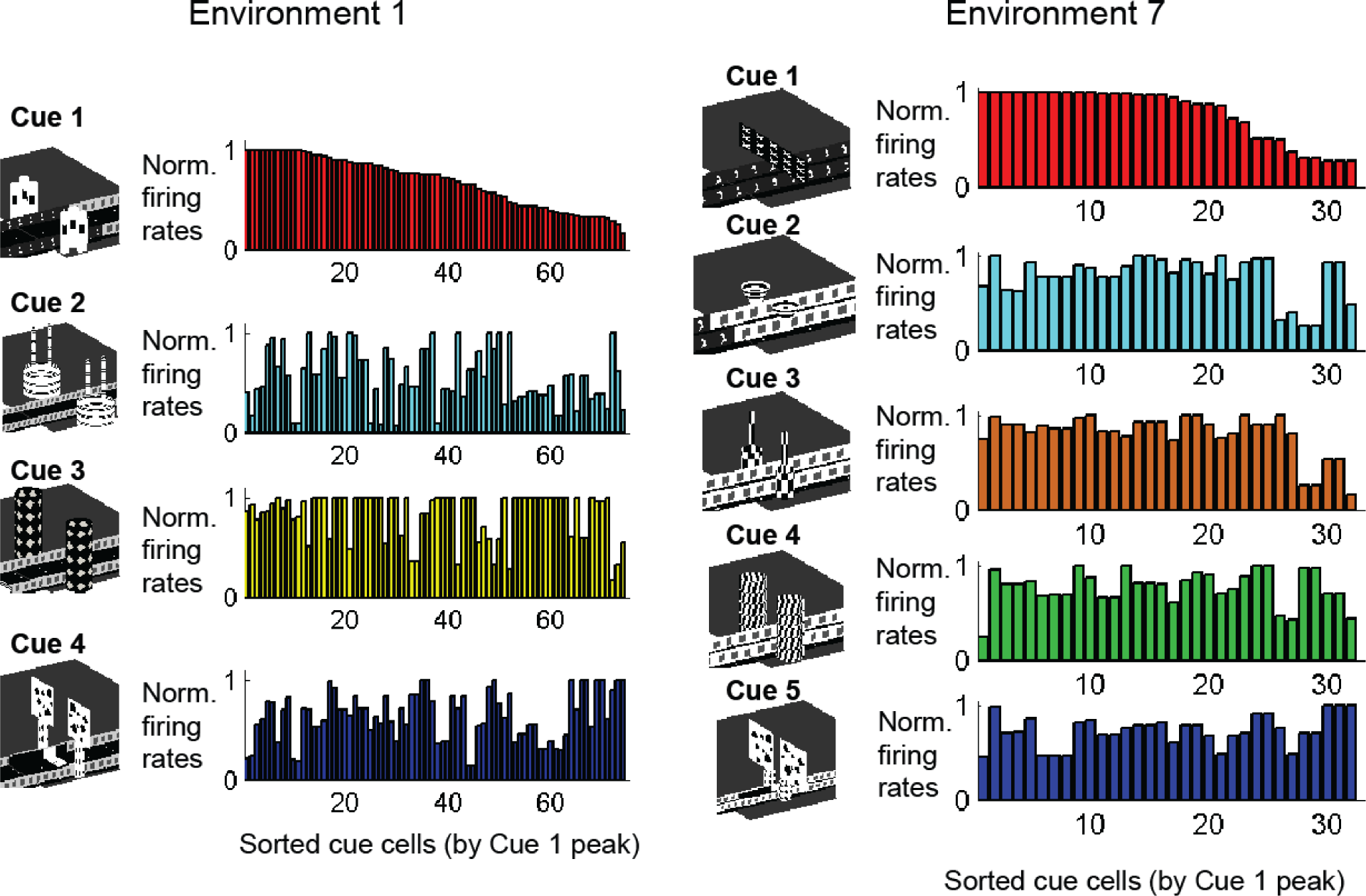
Identity of each cue is encoded by population of cue cells. Cue cell firing rates around individual cues in two different environments (left and right). In each environment around each cue, cue cell firing rates are sorted by the relative peak amplitude of the spatial firing rates at the first cue of the environment. Firing rate peaks for subsequent cues are sorted in the same order as for the first cue. The firing rate peak amplitude for a cue cell at a single cue is not predictive for the amplitude of the peaks at other cues but is predictive for the spatial shifts of the peaks from other cues. For this reason, the amplitude of the firing rate peaks might encode cue features or identity.

## References

Aronov, D., and Tank, D.W. (2014). Engagement of neural circuits underlying 2D spatial navigation in a rodent virtual reality system. Neuron 84, 442-456.

Barry, C., and Burgess, N. (2014). Neural mechanisms of self-location. Curr Biol 24, R330-339.

Boccara, C.N., Sargolini, F., Thoresen, V.H., Solstad, T., Witter, M.P., Moser, E.I., and Moser, M.B. (2010). Grid cells in pre‐ and parasubiculum. Nat Neurosci 13, 987-994.

Brun, V.H., Leutgeb, S., Wu, H.Q., Schwarcz, R., Witter, M.P., Moser, E.I., and Moser, M.B. (2008). Impaired spatial representation in CA1 after lesion of direct input from entorhinal cortex. Neuron 57, 290-302.

Burak, Y., and Fiete, I.R. (2009). Accurate path integration in continuous attractor network models of grid cells. PLoS Comput Biol 5, e1000291.

Bush, D., Barry, C., Manson, D., and Burgess, N. (2015). Using Grid Cells for Navigation. Neuron 87, 507-520.

Calton, J.L., Stackman, R.W., Goodridge, J.P., Archey, W.B., Dudchenko, P.A., and Taube, J.S. (2003). Hippocampal place cell instability after lesions of the head direction cell network. J Neurosci 23, 9719-9731.

Calton, J.L., Turner, C.S., Cyrenne, D.L., Lee, B.R., and Taube, J.S. (2008). Landmark control and updating of self-movement cues are largely maintained in head direction cells after lesions of the posterior parietal cortex. Behav Neurosci 122, 827-840.

Campbell, M.G., Ocko, S.A., Mallory, C.S., Low, I.I.C., Ganguli, S., and Giocomo, L.M. (2018). Principles governing the integration of landmark and self-motion cues in entorhinal cortical codes for navigation. Nat Neurosci 21, 1096-1106.

Carpenter, F., Manson, D., Jeffery, K., Burgess, N., and Barry, C. (2015). Grid cells form a global representation of connected environments. Current biology : CB 25, 1176-1182.

Chen, G., Manson, D., Cacucci, F., and Wills, T.J. (2016). Absence of Visual Input Results in the Disruption of Grid Cell Firing in the Mouse. Curr Biol 26, 2335-2342.

Clark, B.J., Bassett, J.P., Wang, S.S., and Taube, J.S. (2010). Impaired head direction cell representation in the anterodorsal thalamus after lesions of the retrosplenial cortex. J Neurosci 30, 5289-5302.

Clark, B.J., Rice, J.P., Akers, K.G., Candelaria-Cook, F.T., Taube, J.S., and Hamilton, D.A. (2013). Lesions of the dorsal tegmental nuclei disrupt control of navigation by distal landmarks in cued, directional, and place variants of the Morris water task. Behav Neurosci 127, 566-581.

Clark, B.J., Sarma, A., and Taube, J.S. (2009). Head direction cell instability in the anterior dorsal thalamus after lesions of the interpeduncular nucleus. J Neurosci 29, 493-507.

Clark, B.J., and Taube, J.S. (2009). Deficits in landmark navigation and path integration after lesions of the interpeduncular nucleus. Behav Neurosci 123, 490-503.

Dana, H., Chen, T.W., Hu, A., Shields, B.C., Guo, C., Looger, L.L., Kim, D.S., and Svoboda, K. (2014). Thy1-GCaMP6 transgenic mice for neuronal population imaging in vivo. PLoS One 9, e108697.

Derdikman, D., Whitlock, J.R., Tsao, A., Fyhn, M., Hafting, T., Moser, M.B., and Moser, E.I. (2009). Fragmentation of grid cell maps in a multicompartment environment. Nat Neurosci 12, 1325-1332.

Diehl, G.W., Hon, O.J., Leutgeb, S., and Leutgeb, J.K. (2017). Grid and Nongrid Cells in Medial Entorhinal Cortex Represent Spatial Location and Environmental Features with Complementary Coding Schemes. Neuron 94, 83-92 e86.

Dombeck, D.A., Harvey, C.D., Tian, L., Looger, L.L., and Tank, D.W. (2010). Functional imaging of hippocampal place cells at cellular resolution during virtual navigation. Nat Neurosci 13, 1433-1440.

Dombeck, D.A., Khabbaz, A.N., Collman, F., Adelman, T.L., and Tank, D.W. (2007). Imaging large scale neural activity with cellular resolution in awake, mobile mice. Neuron 56, 43-57.

Domnisoru, C., Kinkhabwala, A.A., and Tank, D.W. (2013). Membrane potential dynamics of grid cells. Nature 495, 199-204.

Erskine, L., and Herrera, E. (2014). Connecting the retina to the brain. ASN Neuro 6.

Evans, T., Bicanski, A., Bush, D., and Burgess, N. (2016). How environment and self-motion combine in neural representations of space. J Physiol 594, 6535-6546.

Finkelstein, A., Derdikman, D., Rubin, A., Foerster, J.N., Las, L., and Ulanovsky, N. (2015). Three dimensional head-direction coding in the bat brain. Nature 517, 159-164.

Frohardt, R.J., Bassett, J.P., and Taube, J.S. (2006). Path integration and lesions within the head direction cell circuit: comparison between the roles of the anterodorsal thalamus and dorsal tegmental nucleus. Behav Neurosci 120, 135-149.

Fuhs, M.C., and Touretzky, D.S. (2006). A spin glass model of path integration in rat medial entorhinal cortex. J Neurosci 26, 4266-4276.

Fusi, S., Miller, E.K., and Rigotti, M. (2016). Why neurons mix: high dimensionality for higher cognition. Curr Opin Neurobiol 37, 66-74.

Fyhn, M., Hafting, T., Treves, A., Moser, M.B., and Moser, E.I. (2007). Hippocampal remapping and grid realignment in entorhinal cortex. Nature 446, 190-194.

Gauthier, J.L., and Tank, D.W. (2018). A Dedicated Population for Reward Coding in the Hippocampus. Neuron 99, 179-193 e177.

Geva-Sagiv, M., Las, L., Yovel, Y., and Ulanovsky, N. (2015). Spatial cognition in bats and rats: from sensory acquisition to multiscale maps and navigation. Nat Rev Neurosci 16, 94-108.

Giocomo, L.M. (2016). Environmental boundaries as a mechanism for correcting and anchoring spatial maps. J Physiol 594, 6501-6511.

Giocomo, L.M., Stensola, T., Bonnevie, T., Van Cauter, T., Moser, M.B., and Moser, E.I. (2014). Topography of head direction cells in medial entorhinal cortex. Curr Biol 24, 252-262.

Golob, E.J., and Taube, J.S. (1999). Head direction cells in rats with hippocampal or overlying neocortical lesions: evidence for impaired angular path integration. J Neurosci 19, 7198-7211.

Golob, E.J., Wolk, D.A., and Taube, J.S. (1998). Recordings of postsubiculum head direction cells following lesions of the laterodorsal thalamic nucleus. Brain Res 780, 9-19.

Gu, Y., Lewallen, S., Kinkhabwala, A.A., Domnisoru, C., Yoon, K., Gauthier, J.L., Fiete, I.R., and Tank, D.W. (2018). A Map-like Micro-Organization of Grid Cells in the Medial Entorhinal Cortex. Cell 175, 736-750 e730.

Hafting, T., Fyhn, M., Molden, S., Moser, M.B., and Moser, E.I. (2005). Microstructure of a spatial map in the entorhinal cortex. Nature 436, 801-806.

Hardcastle, K., Ganguli, S., and Giocomo, L.M. (2015). Environmental boundaries as an error correction mechanism for grid cells. Neuron 86, 827-839.

Hardcastle, K., Maheswaranathan, N., Ganguli, S., and Giocomo, L.M. (2017). A Multiplexed, Heterogeneous, and Adaptive Code for Navigation in Medial Entorhinal Cortex. Neuron 94, 375-387 e377.

Harvey, C.D., Coen, P., and Tank, D.W. (2012). Choice-specific sequences in parietal cortex during a virtual-navigation decision task. Nature 484, 62-68.

Harvey, C.D., Collman, F., Dombeck, D.A., and Tank, D.W. (2009). Intracellular dynamics of hippocampal place cells during virtual navigation. Nature 461, 941-946.

Heys, J.G., Rangarajan, K.V., and Dombeck, D.A. (2014). The functional micro-organization of grid cells revealed by cellular-resolution imaging. Neuron 84, 1079-1090.

Hollup, S.A., Kjelstrup, K.G., Hoff, J., Moser, M.B., and Moser, E.I. (2001). Impaired recognition of the goal location during spatial navigation in rats with hippocampal lesions. J Neurosci 21, 4505-4513.

Hopfield, J.J. (2015). Understanding Emergent Dynamics: Using a Collective Activity Coordinate of a Neural Network to Recognize Time-Varying Patterns. Neural Comput 27, 2011-2038.

Hoydal, O.A.S., E. R.; Moser M. B.; Moser E. I. (2018). Object-vector coding in the medial entorhinal cortex. bioRxiv.

Kinkhabwala, A.A., Aronov, D., Tank, D. W. (2015). Visual cue-related activity of MEC cells during navigation in virtual reality In Society for Neuroscience (Chicago).

Kloosterman, F., Davidson, T.J., Gomperts, S.N., Layton, S.P., Hale, G., Nguyen, D.P., and Wilson, M.A. (2009). Micro-drive array for chronic in vivo recording: drive fabrication. J Vis Exp.

Kropff, E., Carmichael, J.E., Moser, M.B., and Moser, E.I. (2015). Speed cells in the medial entorhinal cortex. Nature 523, 419-424.

Krupic, J., Bauza, M., Burton, S., Barry, C., and O’Keefe, J. (2015). Grid cell symmetry is shaped by environmental geometry. Nature 518, 232-235.

Krupic, J., Bauza, M., Burton, S., and O’Keefe, J. (2018). Local transformations of the hippocampal cognitive map. Science 359, 1143-1146.

Larson, B., Abeytunge, S., and Rajadhyaksha, M. (2011). Performance of full-pupil line-scanning reflectance confocal microscopy in human skin and oral mucosa in vivo. Biomed Opt Express 2, 2055-2067.

Lever, C., Burton, S., Jeewajee, A., O’Keefe, J., and Burgess, N. (2009). Boundary vector cells in the subiculum of the hippocampal formation. J Neurosci 29, 9771-9777.

Low, R.J., Gu, Y., and Tank, D.W. (2014). Cellular resolution optical access to brain regions in fissures: imaging medial prefrontal cortex and grid cells in entorhinal cortex. Proc Natl Acad Sci U S A 111, 18739-18744.

Mante, V., Sussillo, D., Shenoy, K.V., and Newsome, W.T. (2013). Context-dependent computation by recurrent dynamics in prefrontal cortex. Nature 503, 78-84.

McNaughton, B.L., Battaglia, F.P., Jensen, O., Moser, E.I., and Moser, M.B. (2006). Path integration and the neural basis of the ‘cognitive map’. Nat Rev Neurosci 7, 663-678.

Mittelstaedt, H.a.M., M.-l. (1982). Homing by path integration (Springer, Berlin, Heidelberg).

Moser, E., Moser, M.B., and Andersen, P. (1993). Spatial learning impairment parallels the magnitude of dorsal hippocampal lesions, but is hardly present following ventral lesions. J Neurosci 13, 3916-3925.

Mukamel, E.A., Nimmerjahn, A., and Schnitzer, M.J. (2009). Automated analysis of cellular signals from large-scale calcium imaging data. Neuron 63, 747-760.

Ocko, S.H., K.; Giocomo, L.; Ganguli, S. (2018). Emergent elasticity in the neural code for space. bioRxiv.

Olsen, G.M., Ohara, S., Iijima, T., and Witter, M.P. (2017). Parahippocampal and retrosplenial connections of rat posterior parietal cortex. Hippocampus 27, 335-358.

Parron, C., Poucet, B., and Save, E. (2004). Entorhinal cortex lesions impair the use of distal but not proximal landmarks during place navigation in the rat. Behav Brain Res 154, 345-352.

Parron, C., and Save, E. (2004). Comparison of the effects of entorhinal and retrosplenial cortical lesions on habituation, reaction to spatial and non-spatial changes during object exploration in the rat. Neurobiol Learn Mem 82, 1-11.

Perez-Escobar, J.A., Kornienko, O., Latuske, P., Kohler, L., and Allen, K. (2016). Visual landmarks sharpen grid cell metric and confer context specificity to neurons of the medial entorhinal cortex. Elife 5.

Pollock, E.D., N.; Wei, X.; Balasubramanian, V. (2018). Dynamic self-organized error-correction of grid cells by border cells. bioRxiv.

Pologruto, T.A., Sabatini, B.L., and Svoboda, K. (2003). ScanImage: flexible software for operating laser scanning microscopes. Biomedical engineering online 2, 13.

Rigotti, M., Barak, O., Warden, M.R., Wang, X.J., Daw, N.D., Miller, E.K., and Fusi, S. (2013). The importance of mixed selectivity in complex cognitive tasks. Nature 497, 585-590.

Rubin, A., Yartsev, M.M., and Ulanovsky, N. (2014). Encoding of head direction by hippocampal place cells in bats. J Neurosci 34, 1067-1080.

Solstad, T., Boccara, C.N., Kropff, E., Moser, M.B., and Moser, E.I. (2008). Representation of geometric borders in the entorhinal cortex. Science 322, 1865-1868.

Stensola, T., Stensola, H., Moser, M.B., and Moser, E.I. (2015). Shearing-induced asymmetry in entorhinal grid cells. Nature 518, 207-212.

Stewart, S., Jeewajee, A., Wills, T.J., Burgess, N., and Lever, C. (2014). Boundary coding in the rat subiculum. Philos Trans R Soc Lond B Biol Sci 369, 20120514.

Taube, J.S., Kesslak, J.P., and Cotman, C.W. (1992). Lesions of the rat postsubiculum impair performance on spatial tasks. Behav Neural Biol 57, 131-143.

Tsoar, A., Nathan, R., Bartan, Y., Vyssotski, A., Dell’Omo, G., and Ulanovsky, N. (2011). Large-scale navigational map in a mammal. Proc Natl Acad Sci U S A 108, E718-724.

Ulanovsky, N., and Moss, C.F. (2011). Dynamics of hippocampal spatial representation in echolocating bats. Hippocampus 21, 150-161.

Whitlock, J.R., Sutherland, R.J., Witter, M.P., Moser, M.B., and Moser, E.I. (2008). Navigating from hippocampus to parietal cortex. Proc Natl Acad Sci U S A 105, 14755-14762.

Yamahachi, H., Moser, M.B., and Moser, E.I. (2013). Map fragmentation in two‐ and three-dimensional environments. Behav Brain Sci 36, 569-571; discussion 571-587.

Yoon, K., Buice, M.A., Barry, C., Hayman, R., Burgess, N., and Fiete, I.R. (2013). Specific evidence of low-dimensional continuous attractor dynamics in grid cells. Nat Neurosci 16, 1077-1084.

Yoon, K., Lewallen, S., Kinkhabwala, A.A., Tank, D.W., and Fiete, I.R. (2016). Grid Cell Responses in 1D Environments Assessed as Slices through a 2D Lattice. Neuron 89, 1086-1099.

